# The role of *N*-glycosylation in spike antigenicity for the SARS-CoV-2 Gamma variant

**DOI:** 10.1101/2023.04.03.535004

**Authors:** Cassandra L. Pegg, Naphak Modhiran, Rhys Parry, Benjamin Liang, Alberto A. Amarilla, Alexander A. Khromykh, Lucy Burr, Paul R. Young, Keith Chappell, Benjamin L. Schulz, Daniel Watterson

## Abstract

The emergence of SARS-CoV-2 variants alters the efficacy of existing immunity towards the viral spike protein, whether acquired from infection or vaccination. Mutations that impact *N*-glycosylation of spike may be particularly important in influencing antigenicity, but their consequences are difficult to predict. Here, we compare the glycosylation profiles and antigenicity of recombinant viral spike of ancestral Wu-1 and the Gamma strain, which has two additional *N*-glycosylation sites due to amino acid substitutions in the N-terminal domain (NTD). We found that a mutation at residue 20 from threonine to asparagine within the NTD caused the loss of NTD-specific antibody binding. Glycan site-occupancy analyses revealed that the mutation resulted in *N*-glycosylation switching to the new sequon at N20 from the native N17 site. Site-specific glycosylation profiles demonstrated distinct glycoform differences between Wu-1, Gamma, and selected NTD variant spike proteins, but these did not affect antibody binding. Finally, we evaluated the specificity of spike proteins against convalescent COVID-19 sera and found reduced cross-reactivity against some mutants, but not Gamma spike compared to Wuhan spike. Our results illustrate the impact of viral divergence on spike glycosylation and SARS-CoV-2 antibody binding profiles.

## Introduction

In December of 2019, a new coronavirus was detected in Wuhan, China which has subsequently been named severe acute respiratory syndrome coronavirus 2 (SARS-CoV-2). SARS-CoV-2, the causative agent of COVID-19, has accounted for more than 651 million infections and more than six million deaths worldwide (World Health Organization Coronavirus 2019 (COVID-19) Dashboard; https://covid19.who.int/, Accessed 03 January 2023). Virus genomic sequences are being generated and shared at an unprecedented rate, with more than one million SARS-CoV-2 sequences available via the Global Initiative on Sharing All Influenza DATA(GISAID), permitting near real-time surveillance of the unfolding pandemics.

The SARS-CoV-2 spike (S) glycoprotein is composed of two subunits: S1, which contains the receptor-binding domain (RBD) responsible for interaction with receptors on host cells, and S2, which mediates membrane fusion and viral entry. The S1 subunit contains two highly immunogenic domains, the N-terminal domain (NTD) and the RBD, which are the major targets of neutralising antibodies. While the RBD binds to the host cell receptor angiotensin-converting enzyme 2 (ACE2), the NTD is proposed to interact with auxiliary receptors including DC-SIGN/L-SIGN^1,2^. Serological analysis of plasma or serum from SARS-CoV-2 infected individuals has revealed only ∼6-20% of circulating antibodies target the NTD compared to ∼65-80% that target the RBD, with the remaining ∼4-20% targeting the S2 subunit^3^. Nonetheless, NTD-targeting mAbs can neutralise SARS-CoV2-2 infection *in vitro* and *in vivo*, suggesting they could be useful for COVID-19 prophylaxis or treatment^3-8^. The NTD-directed monoclonal antibodies identified to-date recognise a glycan-free epitope named the NTD-supersite (residues 14-20, 140-158 and 245-264^3,9^). NTD-directed antibodies are a major selective pressure against the virus, and promote the emergence of NTD escape mutations of SARS-CoV-2 variants^3^.

In late 2020, the SARS-CoV-2 variant of concern (VOC), designated P.1 or Gamma, was first detected in Manaus, Amazonas state, Brazil. This lineage was first detected in four travellers returning to Japan from Amazonas state on 2 January 2021^10^ and was soon recognised as an emergent lineage in Manaus^11^. Gamma evolved from a local B.1.1.28 clade and replaced the parental lineage in <2 months. The Gamma spike protein harbours multiple substitutions: L18F, T20N, P26S, D138Y, R190S, K417T, E484K, N501Y, D614G, H655Y, T1027I and V1176F (**Fig. 1A**). Most of these mutations localise to the NTD and RBD, which are the major targets of neutralising antibodies in convalescent and vaccinated individuals, raising concerns about the efficacy of available vaccines and therapeutic monoclonal antibodies towards this lineage. Some of these mutations had occurred in other VOCs: E484K was shared with B.1.351 (Beta), and N501Y was shared with B.1.1.7 (Alpha) and B.1.351 (Beta). Both these mutations reduce the neutralisation potency of some monoclonal antibodies^12,13^.

**Fig. 1.**
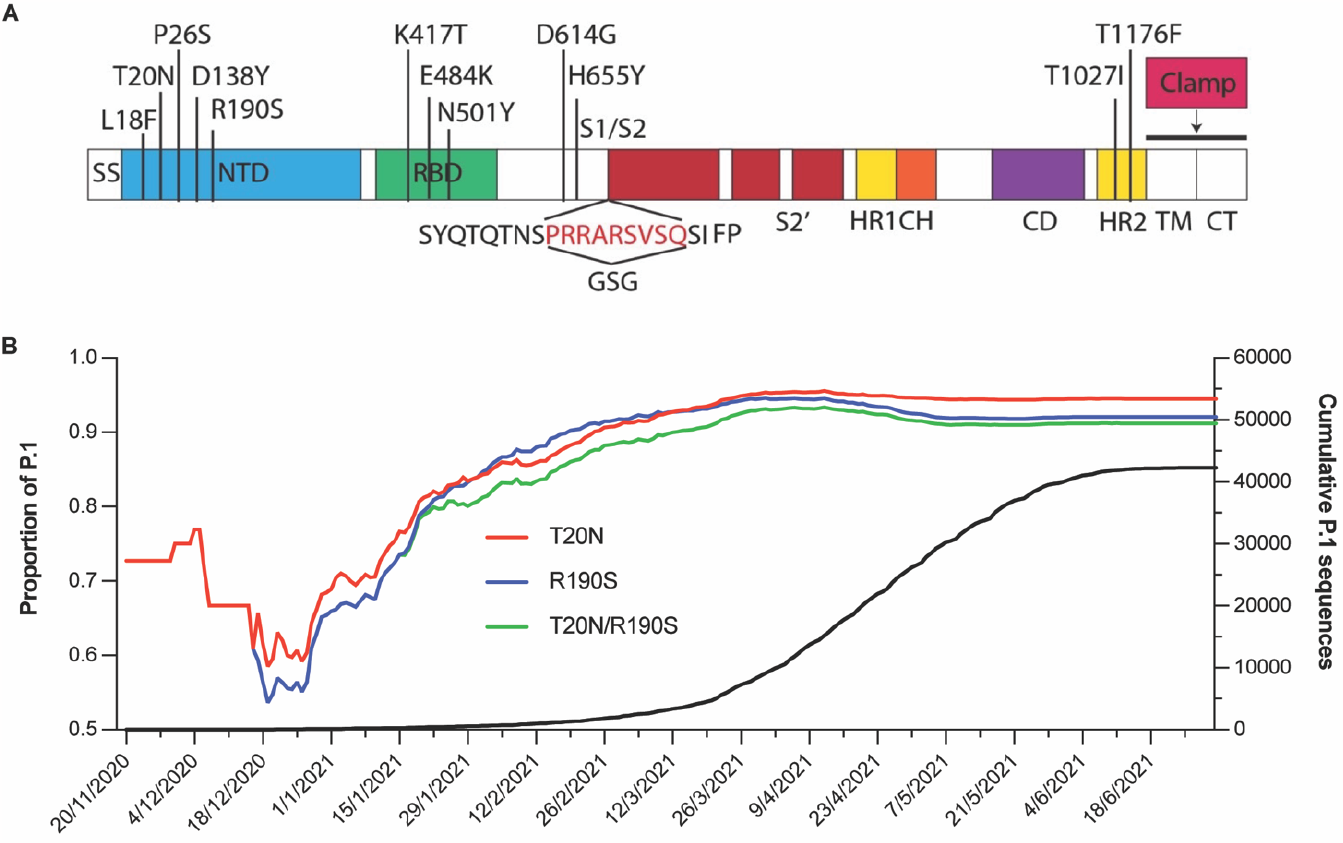
Mutations of the spike protein of the SARS-CoV-2 Gamma variant and evolution of T20N and R190S mutations. **(A)** Schematic of the key domains of spike and locations of the 12 mutations present in the Gamma variant. SS, signal sequence; HR, heptad repeat; CH, central helix; CD, connector domain; TM, transmembrane domain; CT, cytoplasmic domain. The spike construct used in this study comprised residues 1204 of SARS-CoV-2 S with ‘GSG’ substituted at the furin cleavage site and a molecular clamp substituted for the TM and CT domains. **(B)** Early evolution and fixation of SARS-CoV-2 spike glycoforms in Gamma (P.1 and descendent lineages). The left y-axis shows the proportion of sequences with complete collection dates on GISAID harbouring T20N (red), R190S (blue) or both T20N/R190S (green). The right y-axis shows the cumulative number of P.1 sequences deposited on GISAID between 20 Nov 2020 to 1 Jul 2021.

A key post-translational modification that can also modulate the biophysical properties of proteins is glycosylation, whereby carbohydrate moieties (glycans) are attached to nascent proteins during synthesis in the endoplasmic reticulum (ER)^14^. Asparagine- (*N*-) glycans are attached at acceptor asparagine residues within an amino acid consensus motif or sequon (Asn-Xaa-Thr/Ser; Xaa≠Pro). This peptide motif binds with high affinity to the active site of the oligosaccharyltransferase, the enzyme that catalyses *N*-glycosylation in the ER^15^. Glycans play critical roles in virus-host interactions, including stabilising the conformation of viral and host proteins; serving as viral attachment factors, co-receptors, or receptors; promoting structural conformations that facilitate or enhance receptor binding; shielding or presenting underlying viral epitopes; and acting as antigens to direct immune respones^16-18^. Indeed, the spike protein is heavily glycosylated, with 22 occupied *N*-glycosylation sequons present per protomer^19^ and glycosylation has been shown to influence SARS-CoV-2 infection^20^. Host glycans act as ligands, facilitating virus binding and uptake^21,22^ and inhibiting cellular glycosylation blocks SARS-CoV-2 entry^23^. Furthermore, molecular dynamics (MD) studies indicate that spike *N*-glycosylation sites N165 and N234^24^, and in addition N343^25^, stabilise the protein in an open conformation allowing more favourable interactions with the ACE receptor, and that glycan-protein and glycan-glycan interactions form between ACE and spike^26^.

The glycosylation profile of spike has been extensively investigated^27,28^. Minimal differences in glycosylation have been observed between vaccine constructs of trimeric ancestral spike held in a pre-fusion conformation and native (virion derived) spike^29^. Conversely, glycosylation differs substantially between native spike and a monomeric form of the S1 subunit from a non-stabilised protein construct^30^. These studies indicate that the glycosylation profile of spike is relatively stable when proteins are produced in similar cell lines, and overall spike architecture is conserved. However, specific amino acid substitutions in spike can modify site-specific glycosylation. A single substitution D614G (present in the Alpha variant and subsequent VOCs) and mutations specific to the Alpha variant, alter glycosylation profiles at selected sites compared to the ancestral Wuhan-Hu-1 strain^31,32^.

In addition to sequence variations that may promote conformational changes, mutations can also introduce, remove, or change *N*-glycosylation sequons (NxT/S; x≠Pro) which may alter the structure and function of spike. For instance, a mutation (NST_372_>A) in spike of human SARS-CoV-2 results in the loss of an *N*-glycosylation sequon when compared to other related coronaviruses^33^. MD simulations^25^ and binding assays^33^ reveal that the loss of this site increases the binding affinity of the spike RBD to host ACE2 by stabilising the open conformation of the RBD. In VOCs, a mutation at position 19 in the Delta variant abolishes the *N*-glycosylation sequon containing site N17 (NLT_19_>R) but has little impact on the overall glycosylation profile of spike compared to the ancestral strain^34^. The Gamma variant mutations T20N and R190S introduce two new *N*-glycosylation sequons, N_20_RT and NLS_190_, respectively. In addition, the Gamma substitution L18F changes the middle amino acid within the native *N*-glycosylation sequon containing site N17 (NF_18_T). The introduced site N188 in Gamma is occupied with oligomannose type glycans^34,35^, and occupancy of the second introduced site, N20, has been confirmed through cryo-EM mapping^13,35^ and was found to contain predominantly complex type glycans^34,36^. Intriguingly, molecular modelling of the Gamma spike NTD predicts additional N20 and N188 glycans could sterically block NTD-specific antibodies targeting the supersite^35^. To our knowledge, the impact of these Gamma NTD mutations on spike-antibody binding has not been described.

With new and emerging variants and the threat of reduced neutralising antibody capacity, understanding the effects of different mutations on the antibody and glycosylation profiles of the NTD is critical to fully understand SARS-CoV-2 immunity during pandemics. Here, we produced nine trimeric prefusion-stabilised spike proteins that included the ancestral spike (Wu-1), the Gamma variant with all 12 substitutions and Wu-1 spike with single, double or triple mutagenesis of L18F, T20N or R190S. We analysed the mAb binding profiles of each spike protein and measured site-specific *N*-glycosylation occupancy and glycoform abundance at the acquired N20 and N188 *N*-glycosylation sites, and at the native *N*-glycosylation site N17. In addition, we measured NTD-specific IgG titers in sera from COVID-19 convalescent individuals.

## Results and Discussion

### Sensitivity of Gamma spike and its mutations to NTD-specific mAbs

The emergence of the Brazilian VOC (Gamma lineage, P.1) impacted the epidemiological profile of COVID-19 cases due to its higher transmissibility and immune evasion ability^11,37-40^. It emerged after a period of rapid genetic diversification^38^ and accumulated 17 non-synonymous defining mutations, ten of which are in the S gene. Mutations in the RBD at K417T, E484K and N501Y are involved in immune escape^9^, while the NTD also contained five mutations, two of which, T20N and R109S, introduce new *N*-glycosylation sequons (**Fig.1A**). Interestingly, sequence analysis revealed the early detection of mutations at T20N and R190S prior to the description of the gamma variant (**Fig.1B**). Although several studies documented Gamma’s increased transmissibility and immune evasion, there is limited data about the glycosylation profile of Gamma and its impact on antibody binding, particularly within the NTD. Such data may help us better understand immune responses to emerging variants and help mitigate the severe impact of the ongoing pandemic.

To examine potential changes in antibody binding of Gamma spike, we generated recombinant trimeric SARS-CoV-2 spike proteins of the ancestral Wu-1 strain and Gamma with a modified furin cleavage site from Chinese Hamster Ovary (CHO) cells, as described previously^41^ and assayed their binding to defined neutralising mAbs originally isolated from SARS-CoV-2 patients (**Fig. 2**). We found that structurally defined epitope-specific monoclonal antibodies (mAbs) including anti-RBD (B38, CB6, CR3022, 2M10B11 and S309) and anti-NTD (4A8, COVA2-17 and COVA1-22) bound differently to Wu-1 and Gamma spike proteins, consistent with previous studies^42-44^. All class I (B38 and CB6) and class IV (CR3022 and 2M10B11) RBD-specific mAbs exhibited lower binding affinity to Gamma spike than to Wu-1 spike, whereas the class III (S309) mAb showed no difference in binding. NTD-specific mAbs also had lower binding affinity to Gamma spike than to Wu-1 spike: 4A8, recognises the NTD supersite; and COVA1-22, where the binding mode has not yet been determined. Interestingly, the COVA2-17, NTD-specific neutralising antibody, bound strongly to Wu-1 spike, but its binding was completely abolished for Gamma spike. In addition to changes in antibody binding, Gamma spike showed slightly lower affinity to monomeric ACE2 (ACE2 FcM). Influenza hemagglutinin specific C05^45^ and anti-clamp^46^ antibodies served as negative and positive controls for these binding assays, and showed no differences in binding. Together, these data showed substantial changes to antibody binding across the RBD and NTD of Gamma spike.

**Fig. 2.**
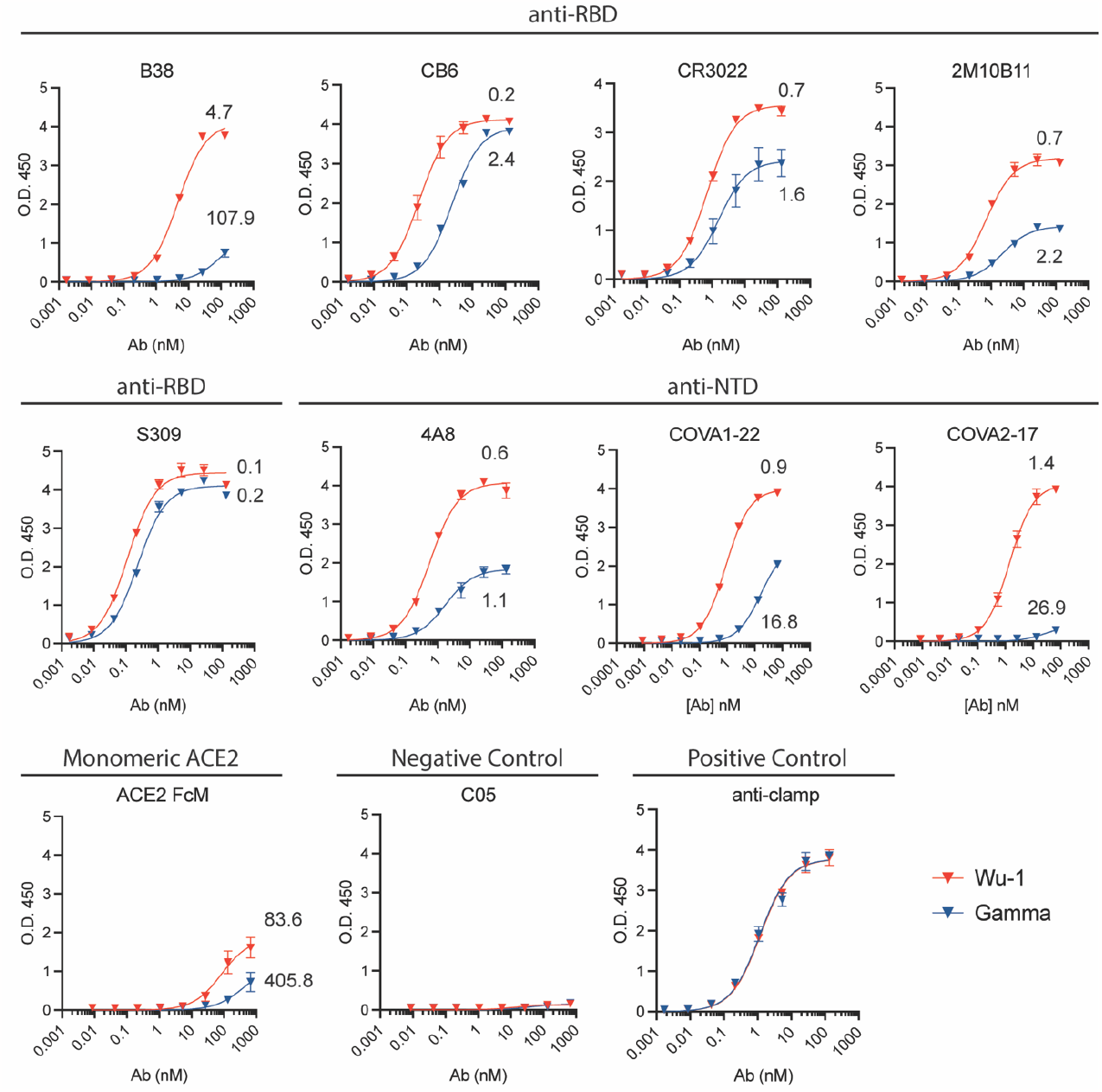
Wu-1 and Gamma SARS-CoV-2 spike proteins exhibit different sensitivities to RBD and NTD specific mAbs. **(A)** Indirect ELISA show binding curves of Wu-1 (Red) and Gamma (Blue) spike to RBD- and NTD-specific antibodies as indicated. Kd shown for each curve, nM.

### Antibody binding of trimeric spike from the Gamma variant with selected mutations

Although multiple antigenic sites, including one at the surface of the NTD, are present on both Wu-1 and Gamma, a single supersite of vulnerability is targeted by neutralising Abs elicited upon infection and vaccination^3,47^. This antigenic supersite (designated site i) comprises the NTD N-terminus (residues 14 to 20), a β-hairpin (residues 140 to 158), and a loop (residues 245 to 264). In the context of this antigenic supersite, the Gamma spike substitutions L18F and T20N are of particular interest, as they alter the native *N*-glycosylation sequon at N17 (NL_18_T > NF_18_T) and introduce a new *N*-glycosylation sequon at N20 (T_20_RT > N_20_RT). Given the proximity of the additional Gamma mutation R190S to the antigenic supersite and predictions that glycans at the resulting introduced *N*-glycosylation sequon (NLS_190_) could alter antibody binding^35^, this substitution is also of high interest. To dissect the consequences of these substitutions, we performed all combinations of single, double or triple mutagenesis at L18F, T20N, and R190S on Wu-1 spike, resulting in seven proteins that were expressed and purified in addition to Wu-1 and Gamma. We then tested binding affinities for selected mAbs to these variant spike proteins (**Fig. 3**). As expected, these substitutions in the NTD did not affect binding of the anti-RBD mAb CB6. Despite anti-NTD mAb COVA1-22 having reduced binding to Gamma, its binding was not reduced to the variant spike proteins with targeted substitutions. In contrast, while anti-NTD COVA2-17 mAb bound Wu-1 spike with high affinity (Kd ∼0.77 nM), the spike variants T20N (Kd ∼8.35 nM), L18F/T20N (Kd >10 nM), and L18F/T20N/R190S (Kd >10 nM) showed substantially reduced or completely abrogated binding for this mAb. COVA2-17 binding affinity did not change with the L18F single mutation (Kd ∼0.5 nM). This data suggests that introduction of a new *N*-glycosylation sequon with the T20N substitution, particularly in combination with L18F, eliminated binding of the anti-NTD COVA2-17 mAb.

**Fig. 3.**
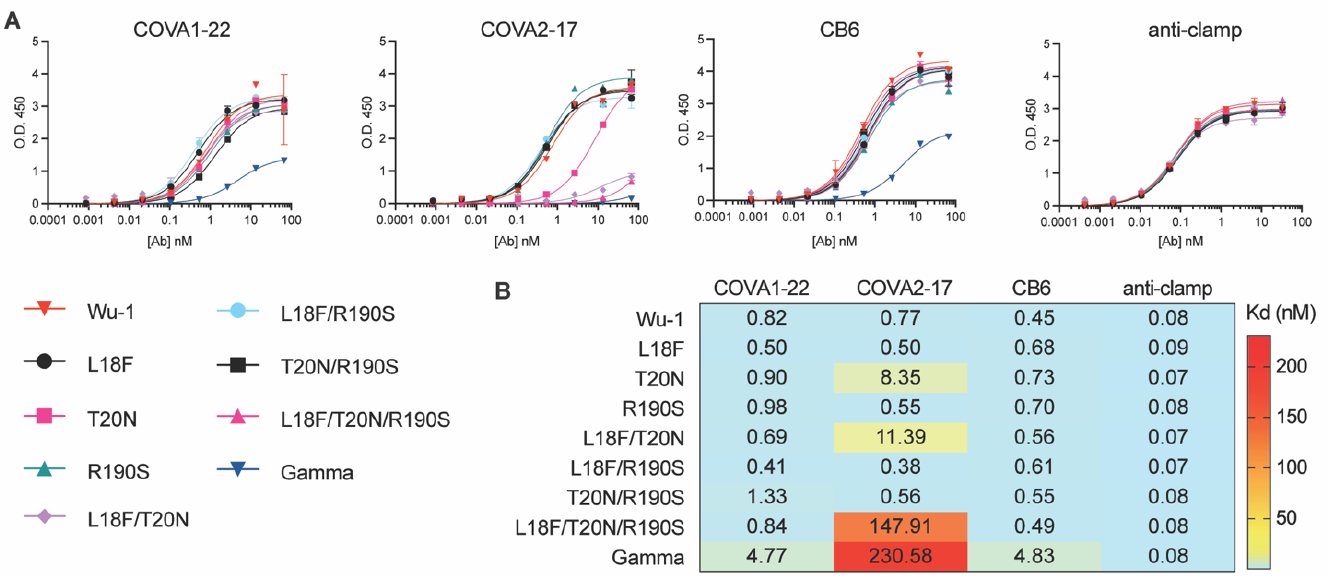
Binding characterisation of SARS-CoV-2 spike proteins of Wu-1, Gamma and NTD mutants. **(A)**. Indirect ELISA show binding curves of RBD- and NTD-specific antibodies as indicated. **(B)** Dissociation constant of mAbs against Wu-1 or Gamma, mutated SARS-CoV-2 spike proteins.

### *N*-glycosylation occupancy of introduced sites N20 and N188 and native N17 in the NTD of Gamma

The introduction of the Gamma mutations T20N and R190S creates two additional *N*-glycosylation sequons (N_20_RT and NLS_190_) in the spike protein, while L18F changes the middle residue of the native N17 sequon proximal to N20 (N_17_F_18_TN_20_RT). To confirm if the decreased antibody binding we observed (**Fig. 3**) was associated with changes in glycosylation at this region, we performed biochemical confirmation of glycosylation site-occupancy in spike from Wu-1, Gamma, and variant spike proteins with selected Gamma substitutions introduced by site-directed mutagenesis. We measured site-specific *N*-glycosylation occupancy and glycoform abundance at the acquired N20 and N188 *N*-glycosylation sites, and at the native *N*-glycosylation site N17. To increase the accuracy with which we could measure site-specific *N*-glycosylation occupancy, we aliquoted pools of peptides and *N*-glycopeptides into two equal portions, one for endoglycosidase treatments with peptide-*N*-Glycosidase F (PNGase F) in ^18^O water to obtain de-glycosylated peptides, and one left untreated for intact glycopeptide analyses. PNGase F removes *N*-glycans converting the previously glycosylated Asn to Asp which results in a mass shift of +0.984 compared to unmodified Asn. In the presence of ^18^O water, the mass difference is increased to +2.984 Da which increases the confidence of site assignment, particularly when multiple *N*-glycosylation sites are potentially present on the same peptide. Using LC-MS/MS, we then measured the abundance of the resulting peptides and de-glycosylated peptides (Supplementary Data 1) and validated *N*-glycosylation occupancy through the identification of glycopeptides (Supplementary Data 2). We observed high occupancy at N17 in the Wu-1 spike. High occupancy was also observed at N17 in Wu-1 spike proteins with one or both of the introduced L18F and R190S mutations (**Fig. 4A**). However, in Gamma spike and all variant proteins with the T20N substitution, the introduction of the sequon at N20 resulted in a switch in site usage, with the new sequon being fully occupied and minimal occupancy at the original N17 site (**Fig. 4A and B**). Similarly, occupancy was high at all variants with the introduced *N*-glycosylation sequon at N188 (**Fig. 4C**). Together, this data showed that loss of binding of the COVA2-17 mAb was associated with *N*-glycosylation site switching from N17 to N20.

**Fig. 4.**
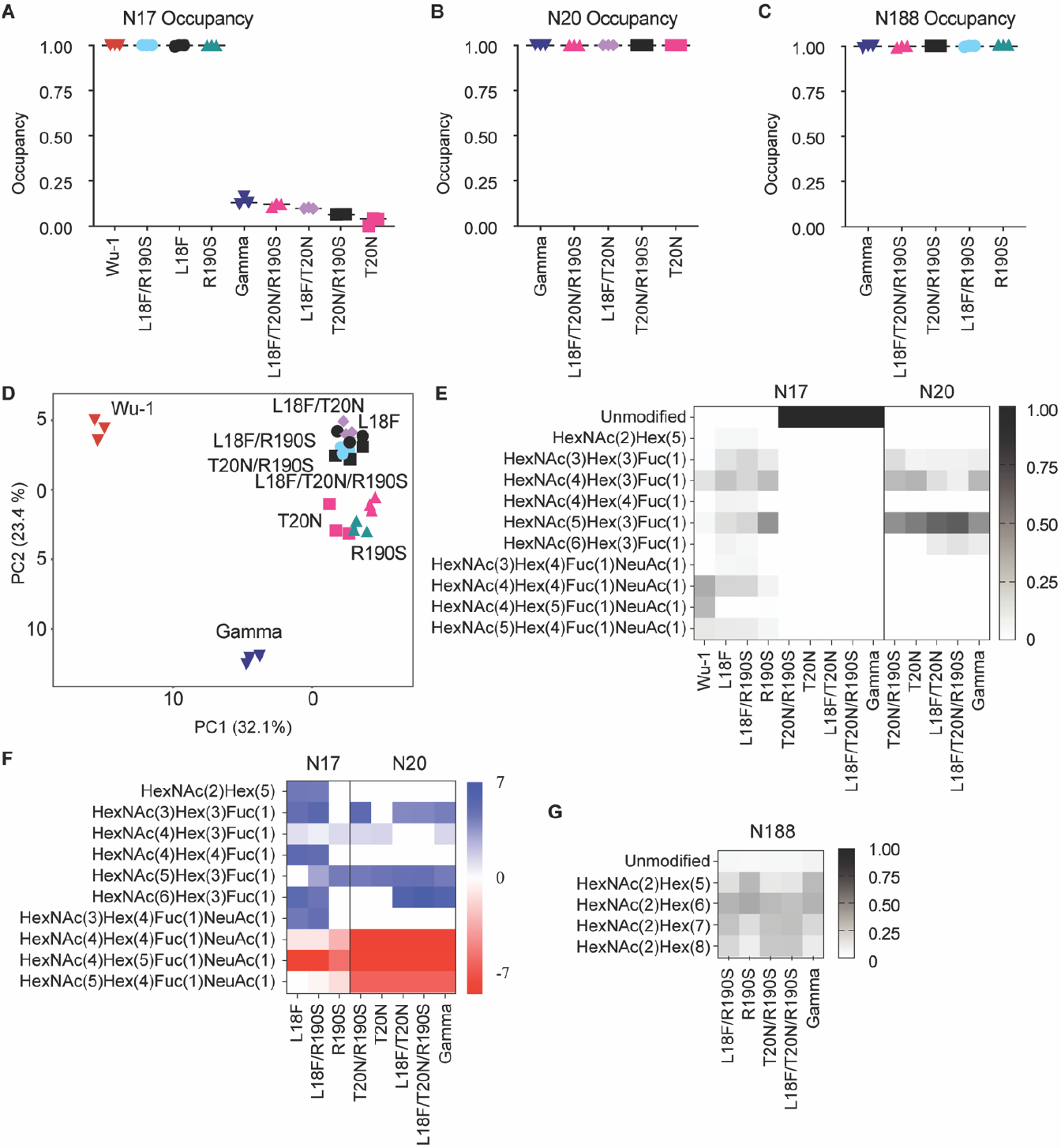
Site-specific *N*-linked glycosylation occupancy and structural heterogeneity of SARS-CoV-2 spike proteins of Wu-1, Gamma and NTD mutants. Site-specific *N-*glycosylation occupancy at sites **(A)** N17, **(B)** N20 and **(C)** N188. **(D)** Principal component analysis (PCA) of glycoform abundances excluding N188. Site-specific glycoform analysis at sites **(E)** N17 and N20 with the **(F)** log_2_ fold change relative to Wu-1 (blue increased abundance, red decreased abundance, P<0.05). For N17 in (E) the unmodified intensity for T20N, L18F/T20N, L18F/T20N/R190S and Gamma were inferred from the occupancy analysis. **(G)** Site-specific glycoform analysis at site N188. HexNAc, *N*-acetylhexosamine; Hex, hexose; Fuc, fucose; NeuAc, *N*-acetylneuraminic acid.

The *N*-linked sites N17 and N20 are adjacent to each other in Gamma (N_17_FTN_20_RT), and glycosylation sites can be skipped when sequons are closely spaced^48^. The competitive higher occupancy at N20 may be due to the type of amino acid in the middle of the *N*-linked sequon (NxS/T; x≠P) as sequons with Leu/L or Phe/F, as at position X of the N17 sequon in the variant proteins, are less efficiently glycosylated^49^.

### Glycoform abundance

The precise glycan structures present at specific *N*-glycosylation sites can be critical for various aspects of spike’s function. We therefore used intact glycopeptide analyses to determine the glycosylation profiles at the native N17, the newly introduced N20 and N188, and six other native *N*-linked sites reliably measured after trypsin digestion (N122, N165, N234, N282, N801, N1098) (Supplementary Data 3 and Supplementary Figure 1). Principal component analysis (PCA) revealed clear separation of glycoforms between Wu-1, Gamma, and the variant spike proteins (**Fig. 4D**). The site-specific glycosylation profile of both N17 and N20 showed predominately complex *N*-glycans with a high degree of fucosylation. These profiles are similar to those reported at N17 after production of trimeric spike in mammalian cell lines^19,26,29,30,34,50,51^, and are consistent with the surface-exposed location of N17 and N20 on the NTD, which would allow processing of glycans to more mature forms^34,35^. Glycan heterogeneity was greater at N17, with more sialylation (NeuAc, *N*-acetylneuraminic acid). The glycans at N20 were more homogenous and contained two predominant forms: HexNAc_4_Hex_3_Fuc_1_ and HexNAc_5_Hex_3_Fuc_1_ (HexNAc, *N*-acetylhexosamine; Hex, hexose; Fuc, fucose) (**Fig. 4E**). Compared to Wu-1, there was a substantial decrease in sialylation in Gamma and all seven variants, with the greatest difference apparent in the variants with the T20N mutation (**Fig. 4F**). Site-specific glycosylation analyses of N188 revealed predominantly oligomannose *N*-glycans for all proteins in which it was present (**Fig. 4G**), in agreement with previous reports^34,35^, and consistent with its inaccessible location within a cleft in the NTD^34^.

To gain insights into the global glycosylation profiles of the various spike proteins, we performed a clustered heatmap analysis of their site-specific *N*-glycosylation profiles (**Fig. 5**). This analysis revealed that Gamma clustered distinctly from Wu-1, while the other variant spike glycoproteins clustered together. This clustering was primarily driven by abundant sialylation in Wu-1, in contrast to abundant fucosylation and short complex glycans (HexNAc_(3-6)_Hex_(3)_) in Gamma. Both Wu-1 and Gamma had similar afucosylation and oligomannose glycans, while these were more abundant in the site-directed mutagenesis variants. To confirm these differences were due to the protein sequence, rather than being expression artefacts, we performed independent expression, purification, and glycosylation analysis, which revealed these glycosylation profiles were robust and reproducible, and were driven by spike protein sequences (Supplementary Figure 1-3 and Supplementary Data 4). Together, our site-specific glycosylation analysis showed that in addition to causing *N*-glycosylation site switching from N17 to N20, the protein sequence changes in Gamma spike caused changes to the *N*-glycan structures at multiple sites across spike, with an overall decrease in sialylation compared to Wu-1.

**Fig. 5.**
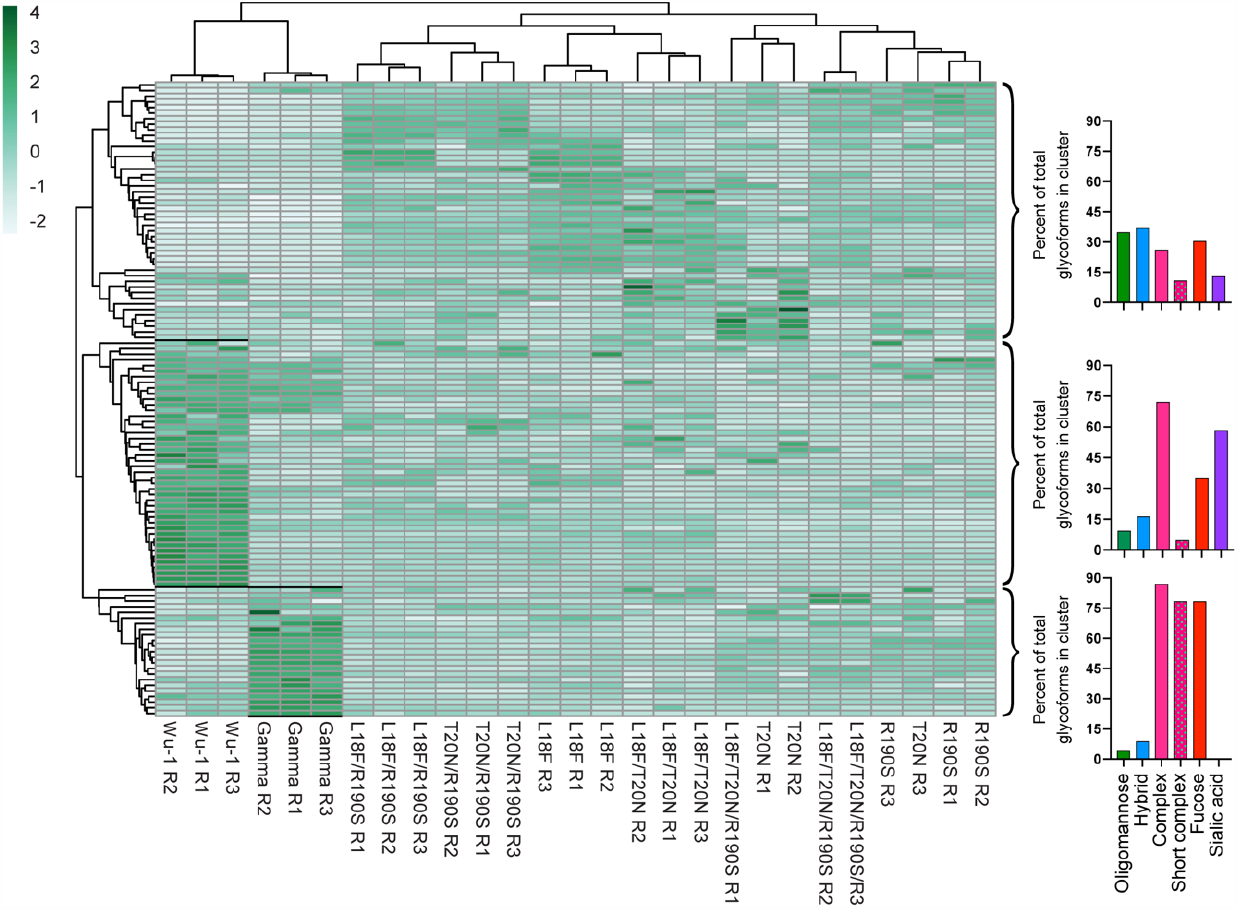
SARS-CoV-2 spike proteins of Wu-1, Gamma and NTD mutants have diverse glycosylation profiles. Clustered heatmap of the relative abundance of all identified glycoforms in all replicates, excluding N188. Both rows and columns were clustered using correlation distance and average linkage. The percentage of types of glycans observed in each selected cluster are represented in bar graphs (Refer to the methods for the allocation of glycan-types). The full clustered heatmap with labelled rows (glycopeptide glycoforms) is shown in Supplementary Figure 3A.

Previous comparisons of the glycosylation of Gamma and Wu-1 spike produced in HEK cells observed less sialylation on Gamma, although less extreme than we observe here in spike produced in CHO cells^34,36^. It is therefore possible that the considerable differences in glycosylation of Gamma and Wu-1 we report here may be cell line specific. Nonetheless, the mechanisms underlying the reduced sialylation of *N*-glycans at diverse sites across Gamma spike compared with Wu-1 spike remain unclear.

### Antibody binding of sialidase treated spike proteins

Our site-specific glycosylation analysis showed two key changes that correlated with reduced binding of select mAbs to Gamma compared to Wu-1 spike: glycosylation site switching from N17 to N20; and lower sialylation at *N*-glycosylation sites across Gamma spike. We therefore tested if this reduced binding was due to *N*-glycosylation site switching or changes in global sialylation. Variant spike proteins were treated with sialidase or left untreated, with removal of sialic acid confirmed by mass spectrometry (Supplementary Data 5). The proteins were then tested for anti-RBD (CB6) and anti-NTD (COVA2-17) binding. No change in mAb binding was observed in any neuraminidase-treated spike compared to untreated spike (**Fig. 6**). These results indicated that the significant changes in sialylation associated with the Gamma mutations were not responsible for the decreased binding of CB6 and COVA2-17 to Gamma, but that these changes in mAb binding were driven by protein sequence changes in the case of CB6, and *N*-glycosylation site switching in the case of COVA2-17.

**Fig. 6.**
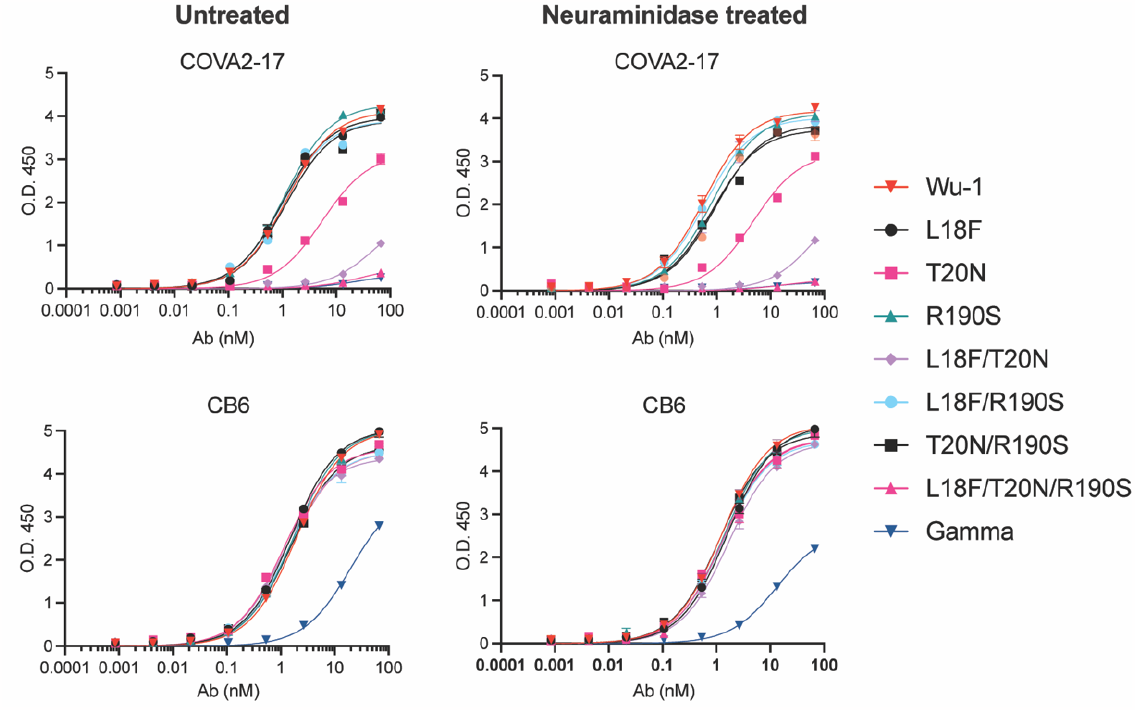
The binding affinities of SARS-CoV-2 spike proteins of Wu-1, Gamma and NTD mutants are not dependant on sialic acid content. Variant spike proteins were treated with neuraminidase for 1 h or left untreated. The proteins were then coated on ELISA plates and binding affinities determined for COVA2-17 or CB6 mAb.

These results are consistent with previous reports showing that global changes in spike glycosylation have minimal influence on its antibody binding profile. For instance, the binding profile of spike to sera from patients previously infected with COVID-19 was similar in spike with near-native glycosylation or which had been engineered to contain only oligomannose glycans^50^. Furthermore, glycosidase digestion of spike glycans to leave only the core GlcNAc elicited effective immune responses in *in vitro* and *in vivo* and protected mice from challenge with the Gamma variant^52^. Glycosylation of spike also likely affects its interactions with innate immune regulators and receptors. For example, spike lacking sialylation has decreased binding avidity for ACE2^52^, while RBDs with complex glycans bind the receptor with greater affinity than RBDs with high mannose glycans^53^. Lectins, including the sialic acid-binding immunoglobulin-like lectin 1 (SIGLEC1) have also been demonstrated to function as attachment factors for SARS-CoV-2^54^. The global changes we observe in Gamma spike glycosylation may therefore have diverse unexpected biophysical or biological consequences.

### ELISA of convalescence sera (3mths after exposure) against NTD domains

The mutations in Gamma spike reduced binding to mAbs with binding sites across the NTD and RBD (**Fig. 2** and **3**). To extend these observations, we characterised the total IgG antibody titers from infected individuals (3 months after infection in 2020, most probably with the Wu-1 strain) against Wu-1, Gamma, and the seven variant spikes. COVID-19 convalescence sera had high affinity to Wu-1, and approximately two to five times lower affinity to L18F, T20N and L18F/T20N/R190S, but not to Gamma (**Fig. 7**). It was unexpected that we observed a reduction in IgG binding titre to spike variants with few targeted site-directed mutations but not in Gamma spike, which contains those same mutations as well as additional sequence changes, including P26S and D138Y. It is possible that the single, double, or triple point mutant variants induce subtle global changes to spike structure or dynamics that perturb domain folding and IgG binding in isolation, but that these changes are rescued by the additional mutations in Gamma Spike.

**Fig. 7.**
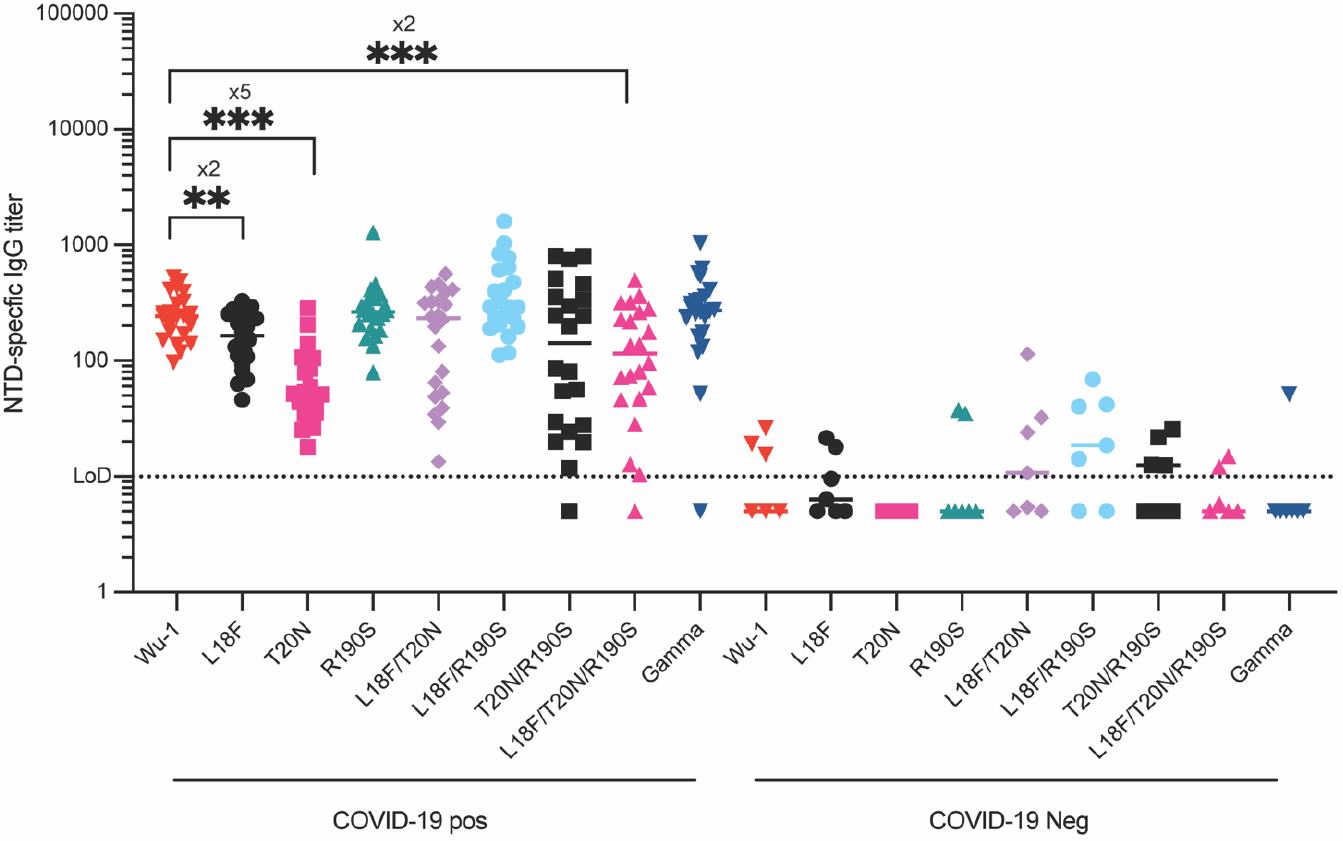
NTD-specific IgG titer of 30 COVID-19 patients (COVID-19 pos) and close contact (COVID-19 neg). Blood samples were obtained from 30 COVID-19 patients and close contact at 3 months after infection IgG titers against the recombinant NTD domains (** *p-value <0*.*01, *** p-value<0*.*001*).

## Conclusion

The emergence of new SARS-CoV-2 variants and evolution of existing variants will result in ongoing diversification of spike protein sequences in circulating virus. Many of these sequence changes may impact a key feature of spike – its glycan shield. The associated impacts of changes in glycosylation on spike function and antibody-mediated immunity are yet to be fully understood. Our results here emphasise that biochemical validation is required to confirm how protein sequence changes that introduce or alter *N*-glycosylation sequons actually affect glycosylation occupancy and associated antibody binding and neutralisation. We found that the structures of *N*-glycans across spike were altered in Gamma compared to Wu-1, highlighting that sequence changes in proteins can globally alter their glycosylation profile. We found no evidence that glycan structure affects antibody binding to spike, but it is likely to impact other spike functions including receptor binding, protein stability, and diverse protein-protein interactions. Our results highlight the importance of continued monitoring of the glycosylation profiles of spike from emerging variants and consideration of the structural, functional, and immunological functions of glycosylation in infection, and in vaccine design and production.

## Methods

### Analysis of P.1 SARS-CoV-2 Spike T20N and R190S variation over time

The metadata of SARS-CoV-2 sequences from the P.1 lineage a with complete collection dates (n=75,879) were downloaded from the GISAID EpiCoV™ portal. Accession IDs with combinations of individual T20N and R190S mutations and combined T20N/R190S mutations were programmatically identified using base Unix grep and sort commands. Cumulative proportions of sequences were calculated using Excel and visualised using GraphPad Prism (v9.0.0).

### Recombinant protein production

For spike protein, soluble, trimeric spikes (residue 1-1204 amino acid) of SARS-CoV 2 /human/China/Wuhan-Hu-1/2019 (GenBank: MN908947), Gamma variant (P.1) (L18F, T20N, P26S, D138Y, R190S, K417T, E484K, N501Y, D614G, D655Y, T1027I and V1176F) were generated using an additional trimerisation domain^41,55^. Mutations at residue 18, (L18F), 20, (T20N) or 190 (R190S) were introduced as single/double/triple mutations by mutagenesis. Spike mutations were added through PCR into the codon-optimised Wuhan reference stain and were cloned into pNBF plasmid via standard infusion cloning procedures. Spike proteins contain substitutions at the furin cleavage site (residues 682–685)^41^. For monoclonal antibodies, heavy and light chains of S309^56^, CB6^57^, B38^58^, CR3022^58^, 2M-10B11^5^, COVA1-22, COVA2-17^59^ and C05^45^ were cloned into a human IgG1 expression vector as described previously^60^. For N-terminal domain, sequence encoding residue 1-305 amino acid of SARS-CoV 2 was cloned in-frame with mannose-binding protein for purification purpose. In brief, proteins were produced in ExpiCHO cells transfected with Expifectamine™ reagent (Thermo Fisher Scientific) as per manufacturer’s protocol^41^. Clarified supernatants were then filtered using 0.22 µm and were affinity purified, concentrated and buffer exchanged into PBS pH7.4.

### Enzyme-linked immunosorbent assay (ELISA)

To test antibodies, SARS-CoV-2 spike variant proteins in PBS pH7.4 were immobilised on Maxisorb ELISA (Nunc) plates at a concentration of 2 μg/mL overnight. Serial 5-fold dilutions of Fc-fusion nanobodies or antibodies in 1X KPL in blocking buffer were incubated with the immobilised antigen, followed by incubation with HRP-coupled anti-human IgG (MilleniumScience) before adding the chromogenic substrate TMB (ThermoFisher). Reactions were stopped with 2 M H_2_SO_4_ and absorption measured at 450 nm.

For sialic acid removal, 15-20 µg of each sample was treated with Neuraminidase A (α2-3,6,8,9 specific, New England Bioloabs, catalogue number P0722S) using 4 U/µg of protein in 100 µL of PBS for 1 h at 37 °C. Confirmation of sialic acid removal was confirmed by mass spectrometry using 2.5-5 µg of protein, which was proteolytically digested as described in *Proteomic and glycoproteomic sample preparation*.

### Proteomic and glycoproteomic sample preparation

Spike protein samples (10 µg of protein) were prepared in triplicate (n=27). Proteins were denatured, reduced, alkylated, quenched and methanol/acetone precipitated overnight as previously described^61^. Protein pellets were resuspended in 50 µL of 50 mM NH_4_HCO_3_ and digested for 16 h with trypsin (Sigma-Aldrich, Product code T6567) with an enzyme to protein ratio of 1:20. The proteolytic enzyme was inactivated by heating for 5 min at 95 °C before incubation with 1 mM phenylmethylsulfonyl fluoride at room temperature for 10 min. The proteolytic digests were aliquoted into two equal volumes (containing 5 µg of protein each, n=54) and one aliquot was dried.

At the same time, 15,000 U of glycerol free PNGase F (New England Bioloabs, catalogue number P0705S) was also dried. From here, all steps were preformed quickly to prevent exposure to air. The freshly dried PNGase F was resuspended in 50 mM NH_4_HCO_3_ prepared in 18O water (Sigma-Aldrich, Product code 329878) and 500 U of PNGase F was added to the dried proteolytic digests. The tubes were flushed with nitrogen and the samples were incubated at 37 °C for 4 h with gentle shaking. The peptides were desalted and concentrated with a C18 ZipTip (10 μL pipette tip with a 0.6 μL resin bed; Millipore, Part No: ZTC18S960) according to the manufacture’s recommendations. Samples were dried and stored at -20 °C then reconstituted in 0.1% formic acid directly before mass spectrometry analysis. Protein samples (5 µg of protein) from a second batch of spike were prepared in triplicate (n=27) on a different day in the same manner described above except that the PNGase F steps were omitted.

### Mass spectrometry for site occupancy analysis

The samples (+/- PNGase F) were randomised and run on an Orbitrap Elite mass spectrometer (Thermo Fisher Scientific) as three full experimental technical replicates (n=54) with ∼100 ng of peptides injected for each chromatographic run using a Dionex UltiMate 3000 uHPLC system (Thermo Fisher Scientific, Bremen, Germany). Solvent A was 1% CH_3_CN in 0.1% (v/v) aqueous formic acid and solvent B was 80% (v/v) CH_3_CN containing 0.1% (v/v) formic acid. Samples were loaded onto a C18 Acclaim™ PepMap™ trap column (100 Å, 5 μm × 0.3 mm × 5 mm, Thermo Fisher Scientific) and washed for 3 min at 30 μL/min before peptides were eluted onto a C18 Acclaim™ PepMap™ column (100 Å, 5 μm × 0.75 mm × 150 mm, Thermo Fisher Scientific) at a flow rate of 0.3 μL/min. Peptides and glycopeptides were separated with a gradient of 3-8% solvent B in 5 min to 50% solvent B over 40 min. Survey scans of peptide precursors from *m/z* of 300 to 1800 were acquired in the Orbitrap at a resolution of 60K (full width at half-maximum, FWHM) at 400 *m/z* using an automatic gain control target of 1,000,000 and maximum injection time of 200 ms. The ten most intense precursors with an intensity over 1,000 and charge states above two were selected for fragmentation by beam-type collision-induced dissociation (CID) using a normalised collision energy of 35% with a precursor isolation window of 2 Da. Fragment ions were acquired in the Orbitrap at a resolution of 30K using an automatic gain control target of 100,000 and maximum injection time of 200 ms.

### Mass spectrometry for glycoform analyses and confirmation of sialic acid removal

For glycoform analyses of each batch, samples were randomised and run as technical replicates with ∼300 ng of peptides injected. Batch 1 (n=27) and 2 (n=27) were run on different days. For confirmation of sialic acid removal, each of the samples (+\- Neuraminidase A) were analysed (n=18). LC-ESI-MS/MS was performed using a Prominence nanoLC system (Shimadzu) coupled to a TripleTof 5600 instrument (SCIEX) using a Nanospray III interface using a data-dependent acquisition method as described^62^ with minor alterations. Peptides and glycopeptides were separated with solvent A (1% CH_3_CN in 0.1% (v/v) aqueous formic acid) and solvent B (80% (v/v) CH_3_CN containing 0.1% (v/v) formic acid) with a gradient of 2-60% solvent B in 45 min. Full MS scans were obtained with a range of *m/z* 350-1800 with accumulation times of 0.5 s. High sensitivity mode was used where the top 20 most intense precursors with charge states of 2-5 and intensities greater than 100 were selected for fragmentation with a collision energy (CE) of 40 V and a 15 V spread. An accumulation time of 0.05 s was used with a scan range of *m/z* 40-1800 and precursors were excluded for 5 s after two selections.

### Data analysis *N*-glycosylation occupancy at N17, N20 and N188

The Sequest HT node in Proteome Discoverer (v. 2.0.0.802 Thermo Fisher Scientific) was used to search RAW files from the PNGase F treated samples. The protein FASTA files contained the relevant SARS-CoV-2 spike-clamp protein sequence without the signal peptide combined with a custom contaminants protein database that included porcine trypsin and PNGase F and the UniProt proteome for Chinese hamster (*Cricetulus griseus*, UP000001075, downloaded 20 March 2018). Cleavage specificity was set to specific, allowing one missed cleavage. Mass tolerances of 10 ppm and 0.02 Da were applied to precursor and fragment ions, respectively. Cys-S-beta-propionamide was set as a static modification and dynamic modifications were set to deamidation of Asn (Asn>Asp, +0.984, and +2.984), pyroglutamic acid formation from N-terminal Gln and mono-oxidised Met. A maximum of four dynamic modifications were allowed per peptide. Confident peptide-to-spectrum matches (PSMs) were assigned using the “Fixed PSM Validator” node and a maximum Delta Cn of 0.05 was applied. Precursor peak areas were calculated using the Precursor Ions Area Detector node. The results from the Sequest HT search were investigated using a modified version of an in-house Python script ^63^. Each Asn residue in an *N*-glycosylation consensus site from identified PSMs (with AUC values greater than 1×10^6) was assigned U (unmodified), D (deamidated), ^18^O-D (deamidated). Occupancy was defined as the proportion of the ion intensity peak area of a peptide with U or D (unmodified) or ^18^O-D (occupied) identified in all charge states to the sum of the intensities of all peptide ions. All PSMs were manually validated.

### Data analysis glycoforms

Searches of the proteolytic digests without PNGase F were used to investigate site-specific glycosylation. Glycopeptide identification was performed using Byonic (v2.13.2, Protein Metrics). Cleavage specificity was fully specific with two missed cleavages allowed. Mass tolerances of 50 ppm and 75 ppm were applied to precursor and fragment ions, respectively. For the samples acquired on an Elite Orbitrap instrument to validate occupancy data, mass tolerances of 10 ppm and 20 ppm were applied to precursor and fragment ions, respectively. Cys-S-beta-propionamide was set as a static modification and dynamic modifications were set to deamidation of Asn (common 2), pyroglutamic acid formation from N-terminal Gln (common 1) and mono-oxidised Met (common 1). The *N*-linked database (rare 1) contained 243 *N*-glycans (Supplementary Table **1**), the *O*-linked database (rare 1) was “6 most common” (Supplementary Table **2**) with the addition of three NeuGc glycans - HexNAc(1)Hex(1)NeuAc(1)NeuGc(1), HexNAc(1)Hex(1)NeuGc(1) and HexNAc(1)Hex(1)NeuGc(2). A maximum of three common modifications and one rare modification were allowed per peptide. Before the glycopeptide searches were conducted, a focused protein database was created in Byonic using the protein FASTA files described in the Sequest HT searches, and the parameters described above but without the dynamic glycan modifications. Glycopeptide PSMs were manually validated and occupancy was calculated using the Molecule Interface of Skyline (v20.2.0.343). A unique list of glycoforms was produced using the Byonic results. The unique list of glycopeptides contained the precursor name, precursor (*m/z*), charge and retention time were inserted into the Transition List in Skyline. A retention time window of 4 min was applied and quantification was performed at MS1 level with a mass tolerance of 0.05 (m/z). Instrument settings were specific to an ABSCIEX 5600 instrument. The results from the Skyline were converted to a readable format for GlypNirO^64^. Heatmaps and bar graphs were produced using PRISM v9.1.0 (GraphPad Software, La Jolla California USA). Principle component analysis (PCA) and clustered heatmaps were produced with ClustVis^65^ where glycoform abundances were normalised to the summed abundance of all detected forms with the same site. Glycans were categorised as follows: Oligomannose, contains HexNAc(2); Hybrid, contains HexNAc(3); Complex, contains more than 3 HexNAc; Complex short, contains Hex(3) and more than 3 HexNAc; Fucose, contains at least one Fuc; Sialic acid, contains at least one NeuAc.

### Human convalescent sera

Subjects were identified from samples stored in the David Serisiser Research Biobank (DSRB) (HREC/14/QPAH/275) at Mater Misericordiae Ltd. Previously biobanked sera from a total of 30 subjects who had been infected with SARS-CoV-2, and 6 non-infected close contacts were used in this project. Subjects were at least 3 months following a PCR defined SARS-CoV-2 infection and were not unwell at the time of sample collection. Samples were predominantly collected during mid to late 2020 and likely represent infection with the original “Wuhan” strain. In addition to sample collection, demographic data and clinical details pertaining to the original infection were also noted. All data and biological samples held within the DSRB are de-identified prior to sharing between collaborating sites. Access to samples for this project was approved by the DSRB biobank advisory committee under project number HREC/MML/68320.

### Human ethics statement

All human sera collections were approved by the relevant ethics committees either by Mater Misericordiae Ltd Human Research Ethics Committee (Reference: HREC/MML/68320) or the University of Queensland (Reference: 2021/HE000139)

## Acknowledgments

We thank Dr Amanda Nouwens and Peter Josh at The University of Queensland, School of Chemistry and Molecular Biosciences Mass Spectrometry Facility for their assistance and expertise.

## Author contributions

C.P. and N.M. conceived the project (C.P. performed glycoproteomic analyses. N.M. performed antigen design and ELISA). R.P. performed sequence analysis. B.L. assisted in protein preparations. A.A.A assisted in ELISA using human sera. L.B. provided resources. A.A.K., L.B., P.R.Y. and K.C. provided intellectual input. C.P., N.M, B.S. and D.W. wrote the manuscript. B.S. and D.W. supervised the project. All authors discussed results and edited the final manuscript. Funding acquisition; D.W., B.S., K.C., P.R.Y. and A.A.K. This work was supported by a NHMRC MRFF Coronavirus Research Response grant APP1202445 to D.W., K.C. and P.R.Y., a NHMRC Ideas Grant APP1186699 to B.S. and a NHMRC Ideas Grant 2012883 to A.A.K.

## Data availability

All genome sequences and associated metadata used in this work are published in GISAID’s EpiCoV database under GISAID Identifier: EPI_SET_230311as. To view the contributors of each individual sequence with details such as accession number, Virus name, Collection date, Originating Lab and Submitting Lab and the list of Authors, visit 10.55876/gis8.230311as The *N*-occupancy and glycoproteomic datasets generated and analysed during the current study are available in the MassIVE repository, [ftp://MSV000091533@massive.ucsd.edu; Username for web access: MSV000091533_reviewer; Password: Gamma_Glyco]

## Supplementary information

Supplementary Figures 1-3

Supplementary Data 1_N-linked Occupancy

Supplementary Data 2_Occupancy Validation

Supplementary Data 3_Batch 1 Glycoform Analysis

Supplementary Data 4_ Batch 2 Glycoform Analysis

Supplementary Data 5_Sialidase Glycoform Analysis

