## Supplementary Figures 1-3 for "The role of *N*-glycosylation in spike antigenicity for the SARS-CoV-2 Gamma variant"

**Supplementary Figure 1**. **Site-specific glycoform analysis of two batches of SARS-CoV-2 Spike**. For each *N*-linked site a heatmap of differentially abundant glycoforms is shown with the $\log_2$ fold change relative to Wu-1 (blue increased abundance, red decreased abundance, P<0.05) along with a heatmap of the relative abundance of glycoforms (HexNAc, *N*-acetylhexosamine; Hex, hexose; Fuc, fucose; NeuAc, *N*-acetylneuraminic acid). The proteins names 18, 20 and 190 represent the proteins with the mutations L18F, T20N and R190S, respectively. A heat map of the $\log_2$ fold change relative to Wu-1 is not provided for N188 as the site is not present in Wu-1.

### A. Sites N17 and N20 Batch 1

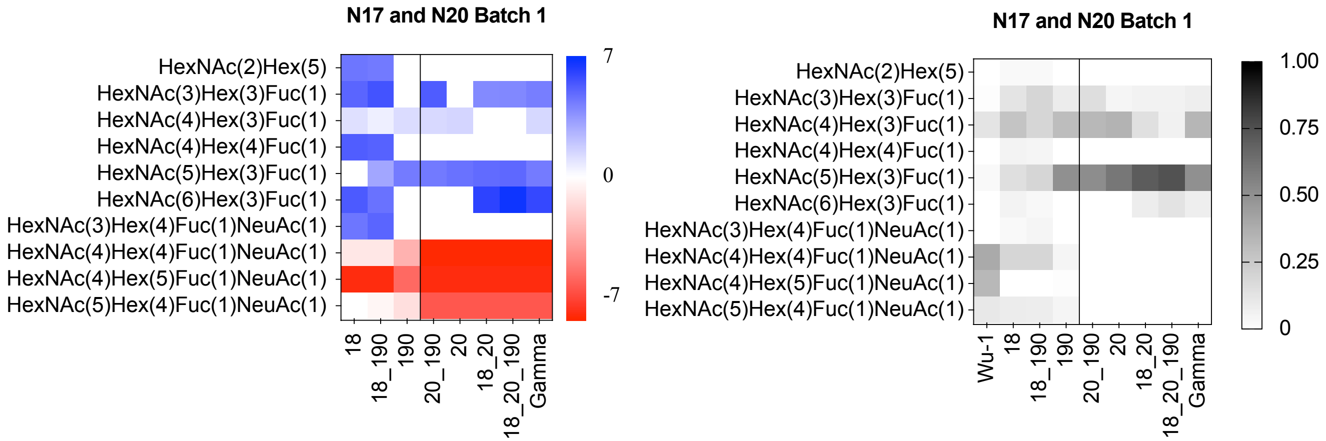

B. Sites N17 and N20 Batch 2

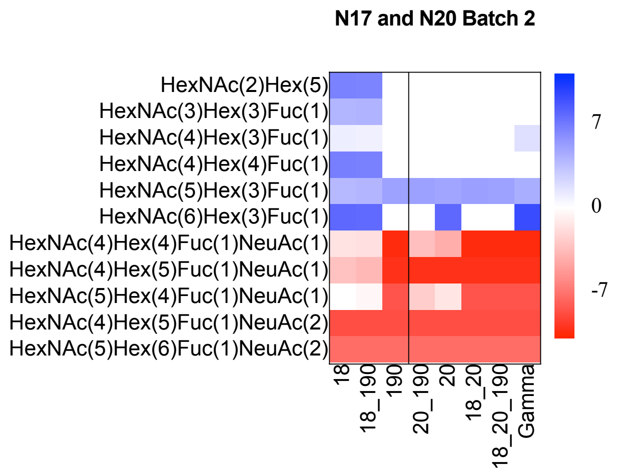


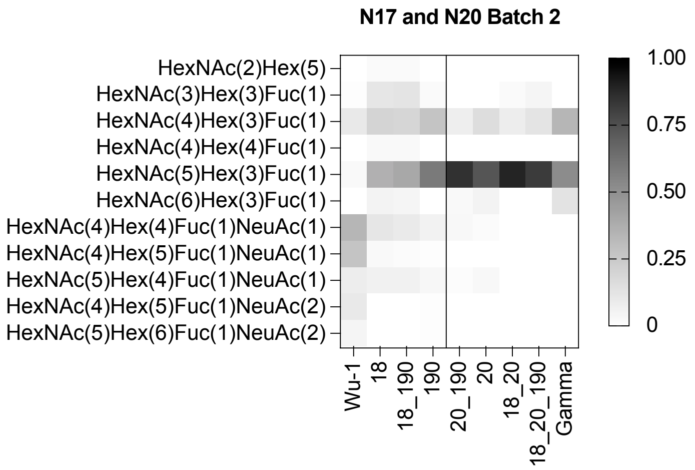

**C. Site N188 Batch 1**

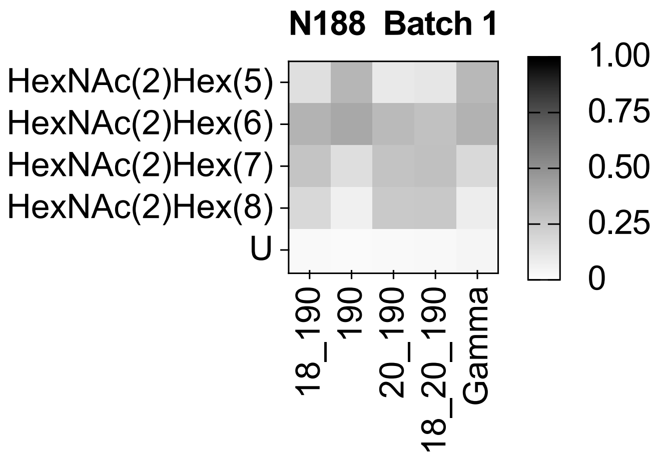


**D. Site N188 Batch 2**

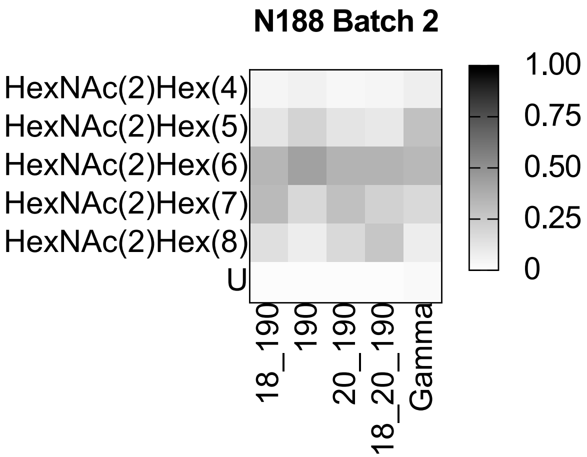

### E. Site N122 Batch 1

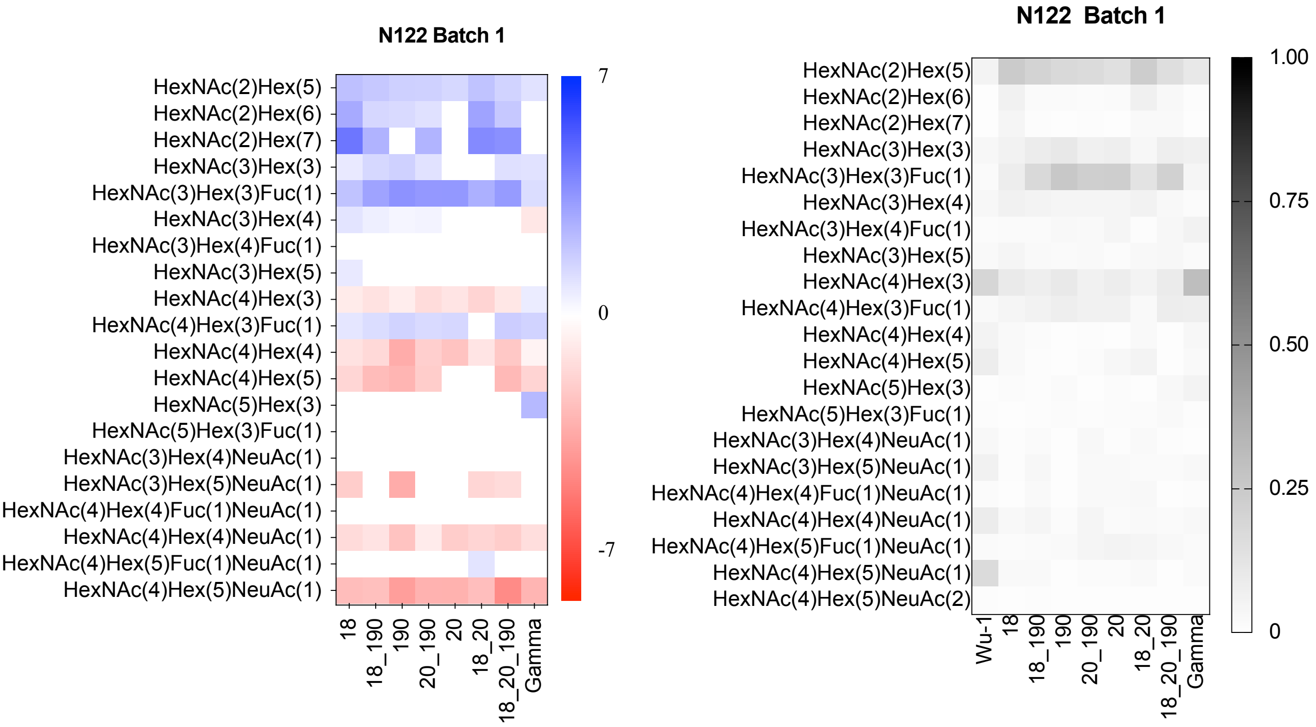


### F. Site N122 Batch 2

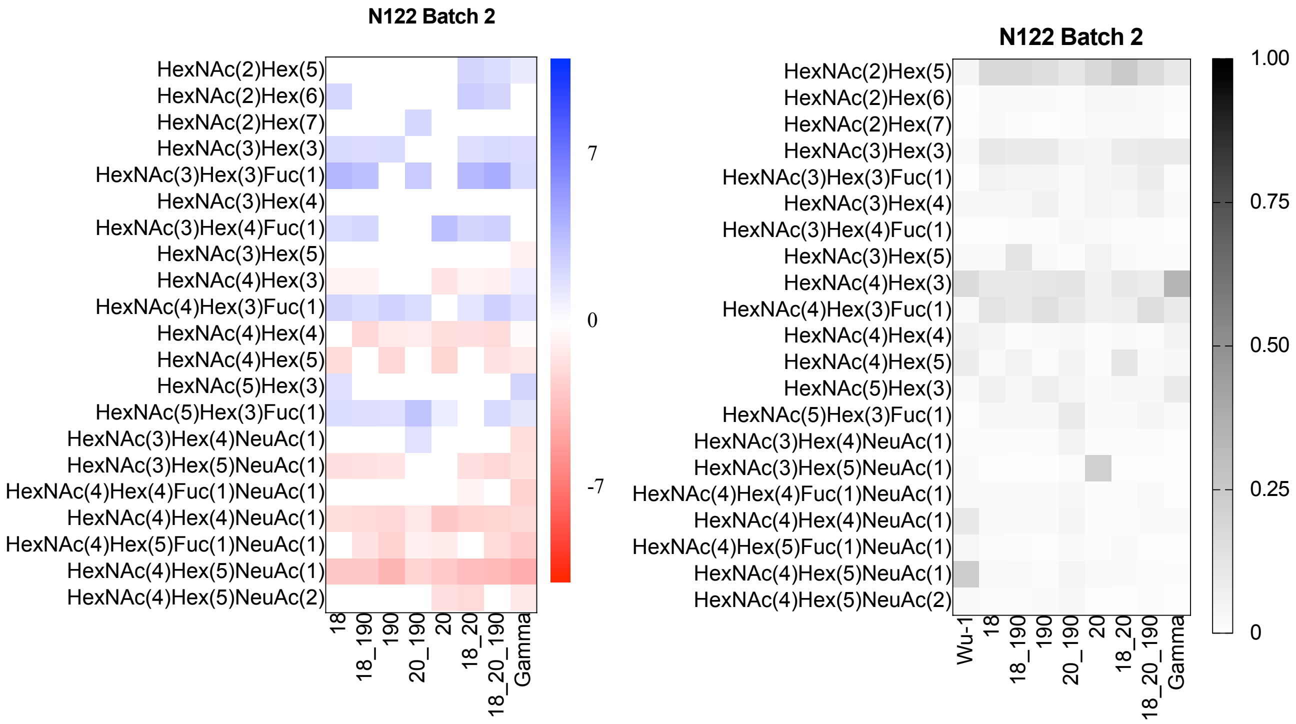

### G. Site N165 Batch 1

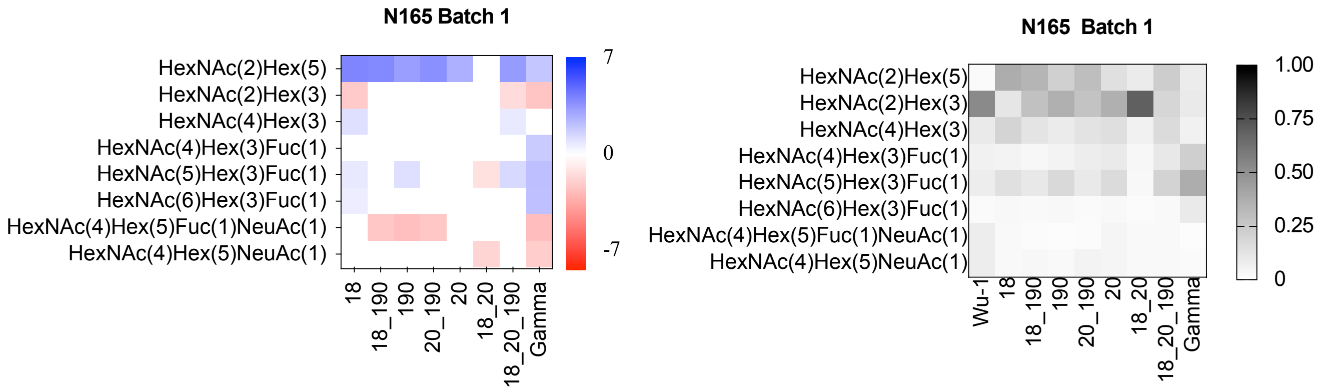


### H. Site N165 Batch 2

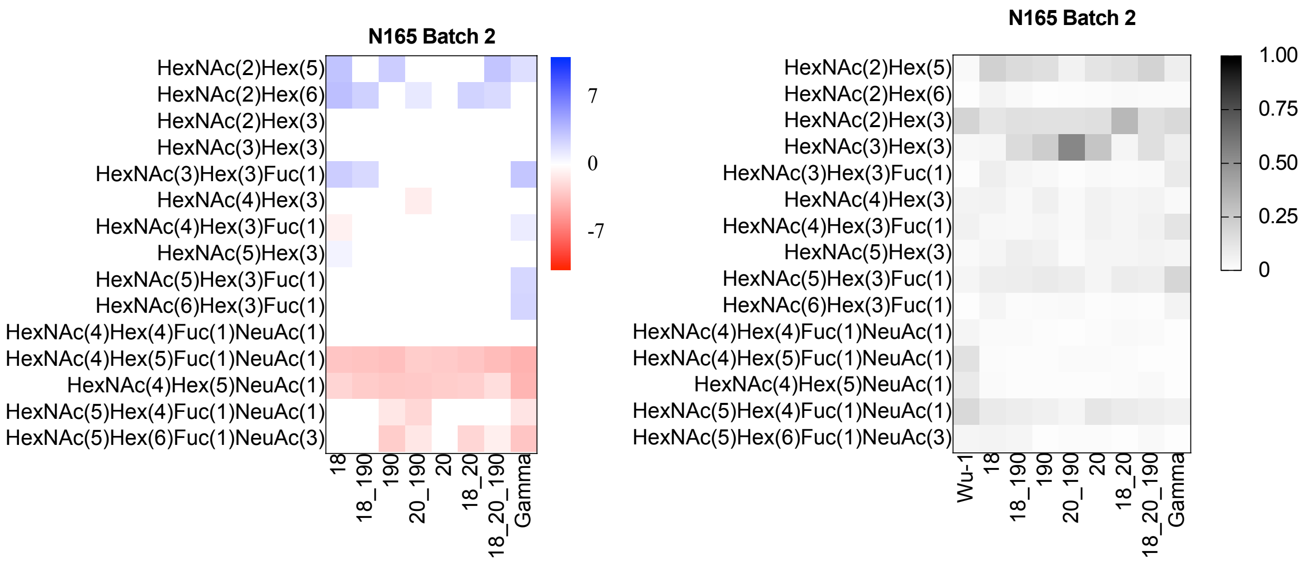

### I. Site N234 Batch 1

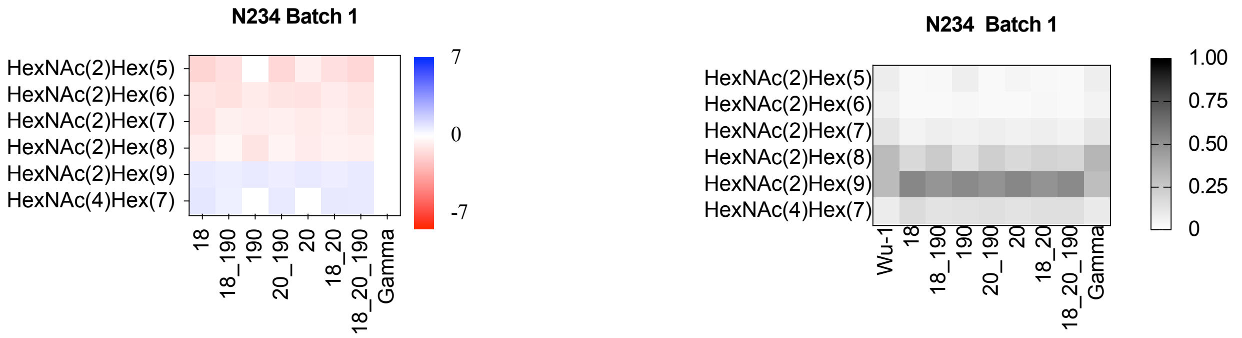


### J. Site N234 Batch 2

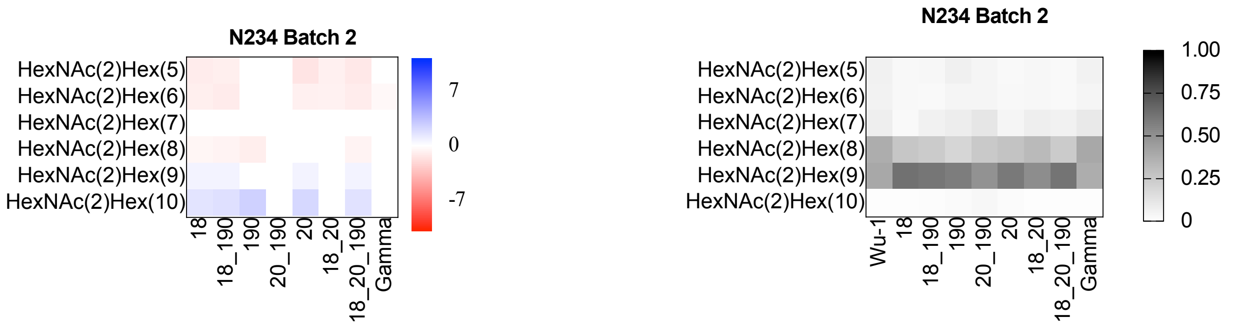

### K. Site N282 Batch 1

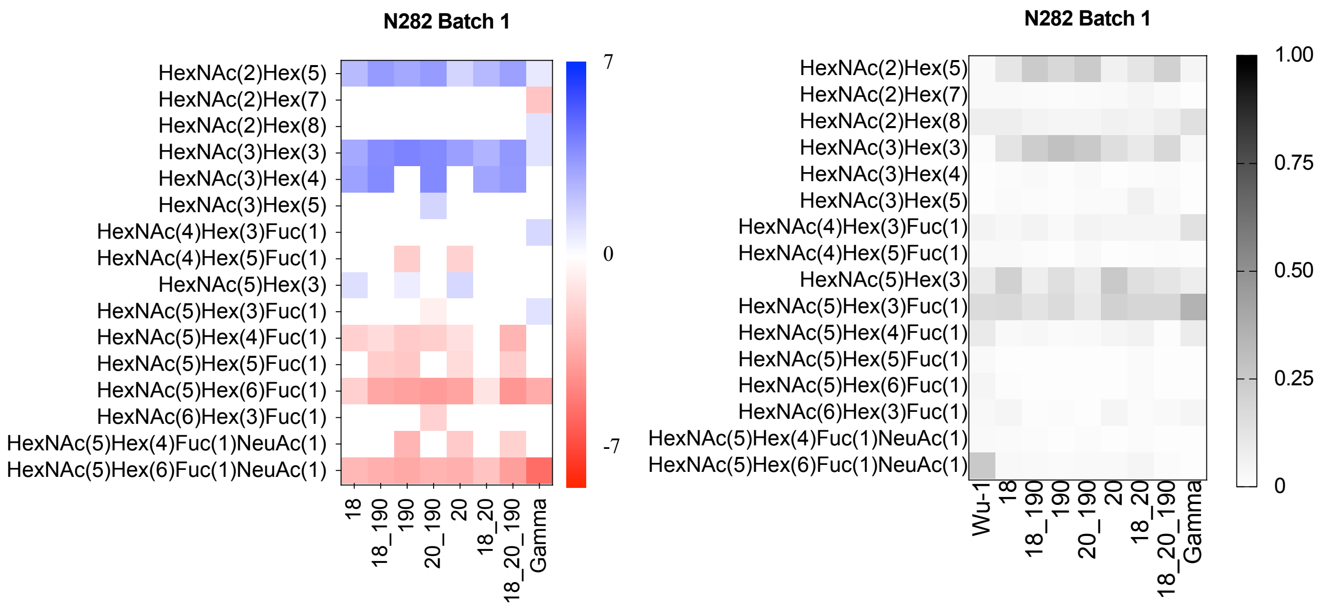


### L. Site N282 Batch 2

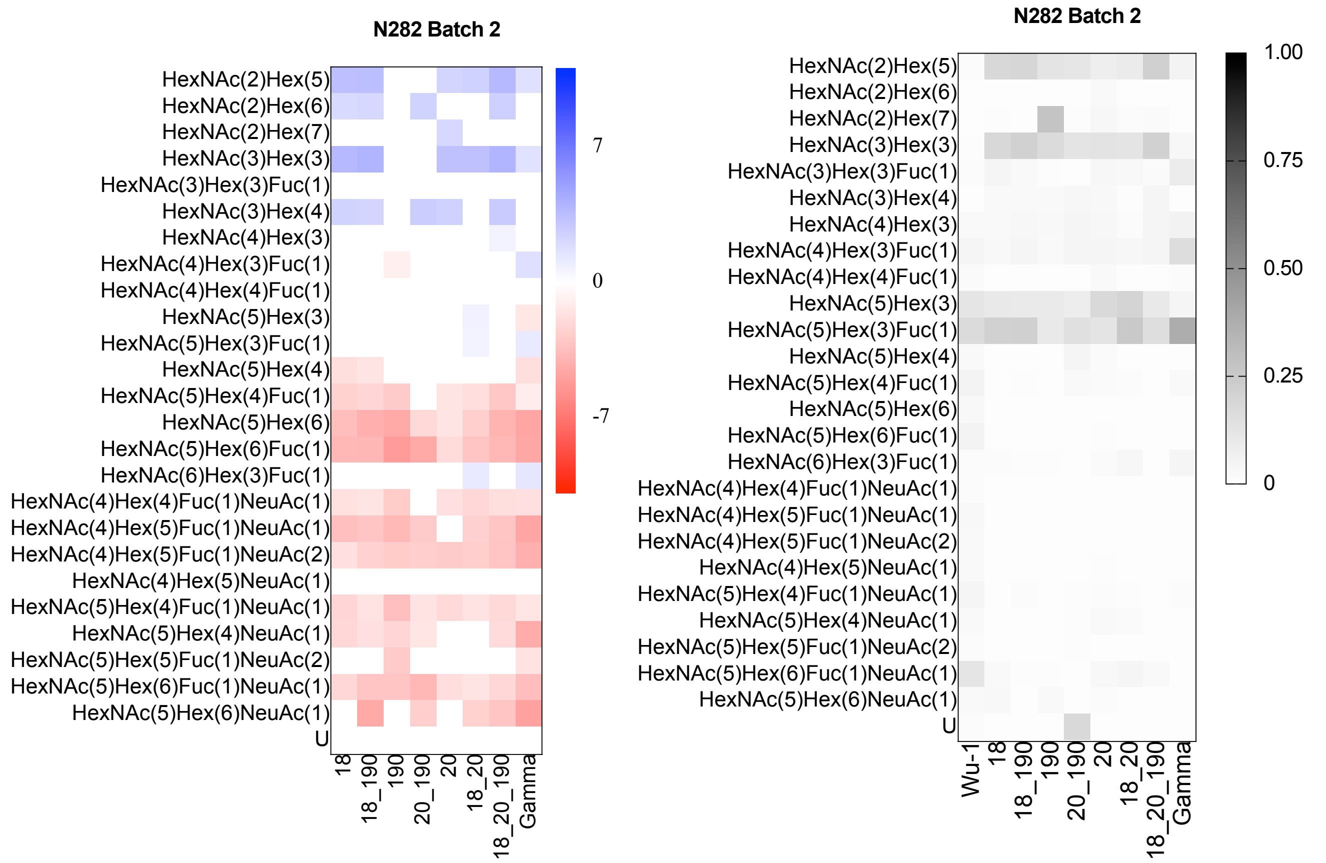

### M. Site N801 Batch 1

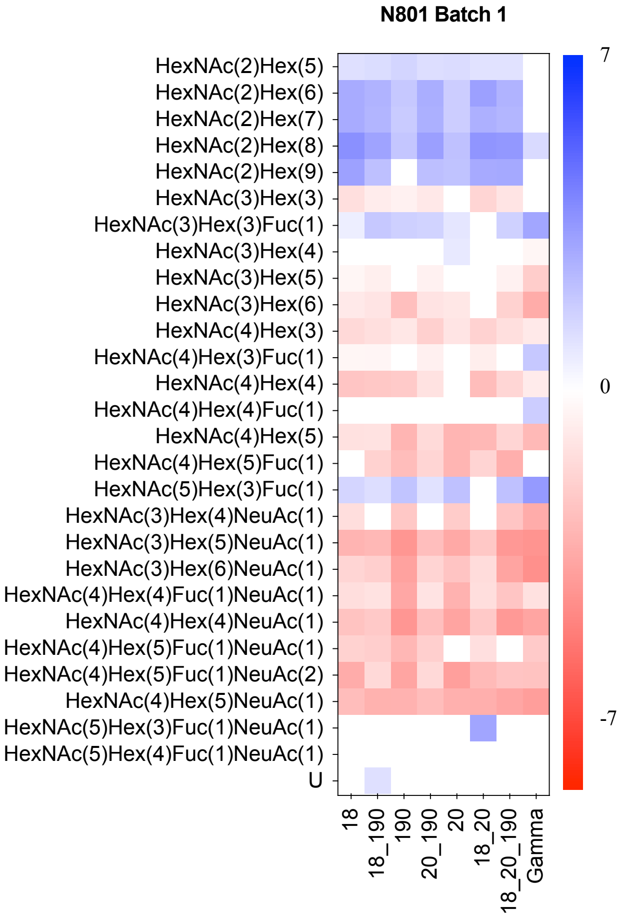


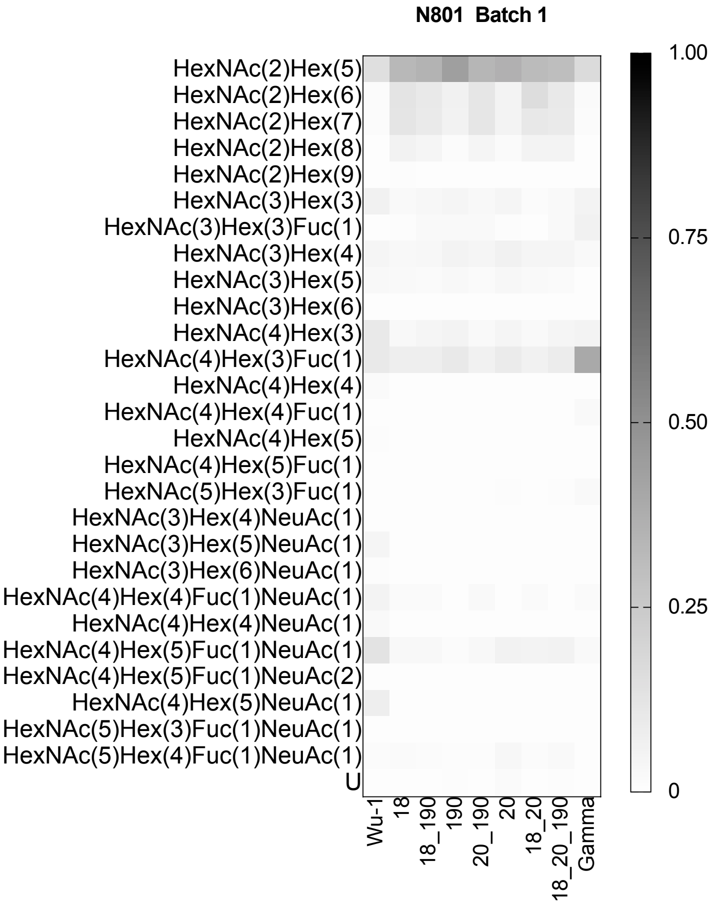


### N. Site N801 Batch 2

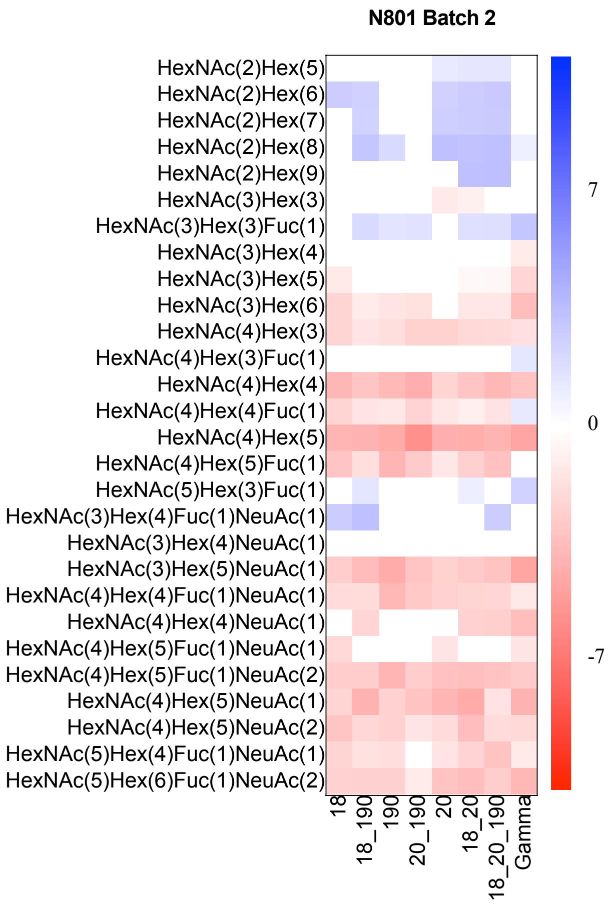


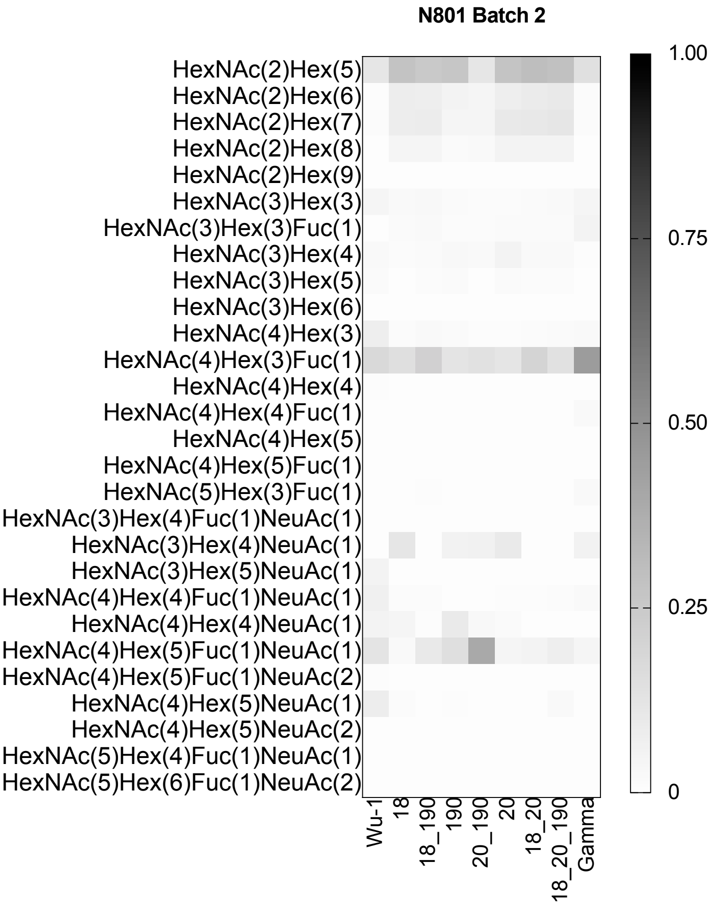

### O. Site N1098 Batch 1

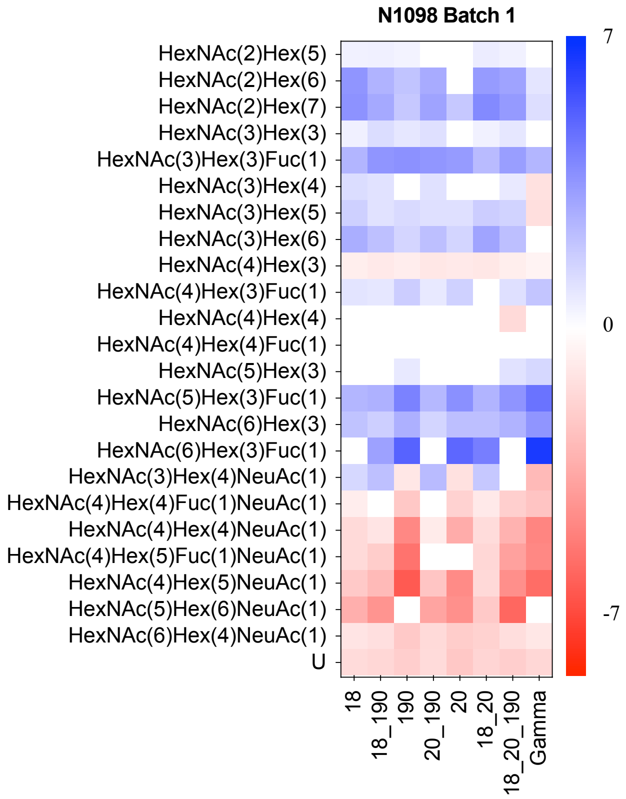


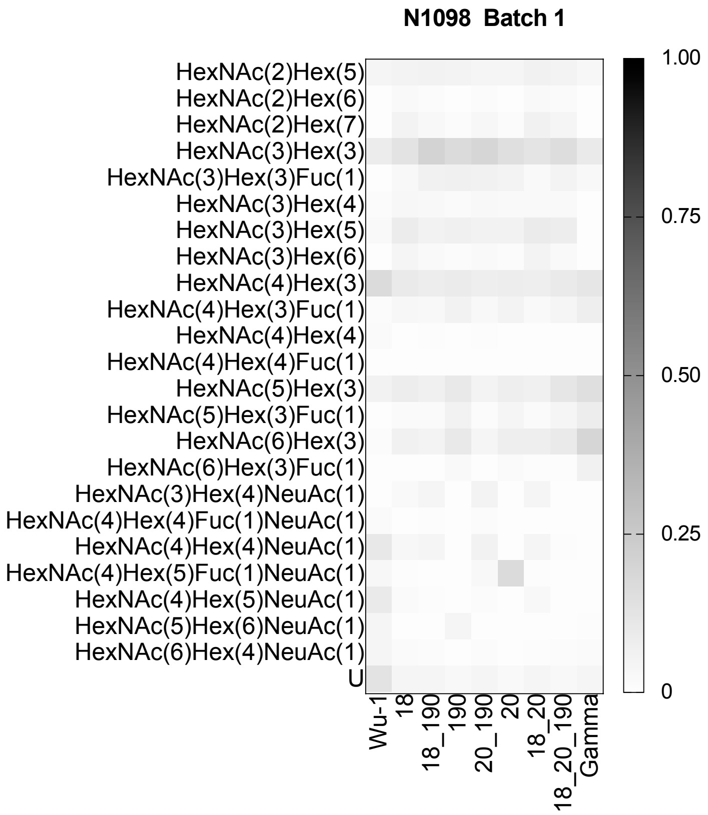


### P. Site N1098 Batch 2

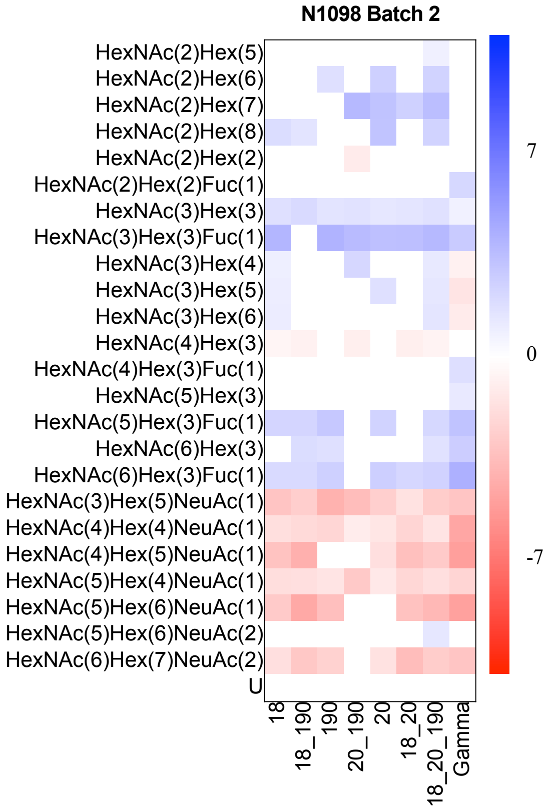


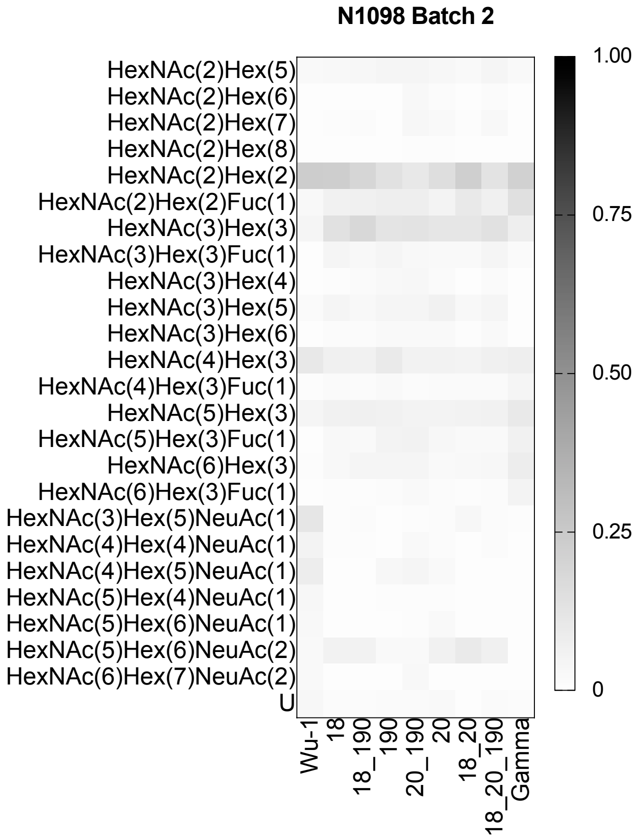

**Supplementary Figure 2**. Principal component analysis of all reliably measured *N*-linked glycoform abundances excluding N188. The proteins names 18, 20 and 190 represent the proteins with the mutations L18F, T20N and R190S, respectively.

**A. Batch 1**

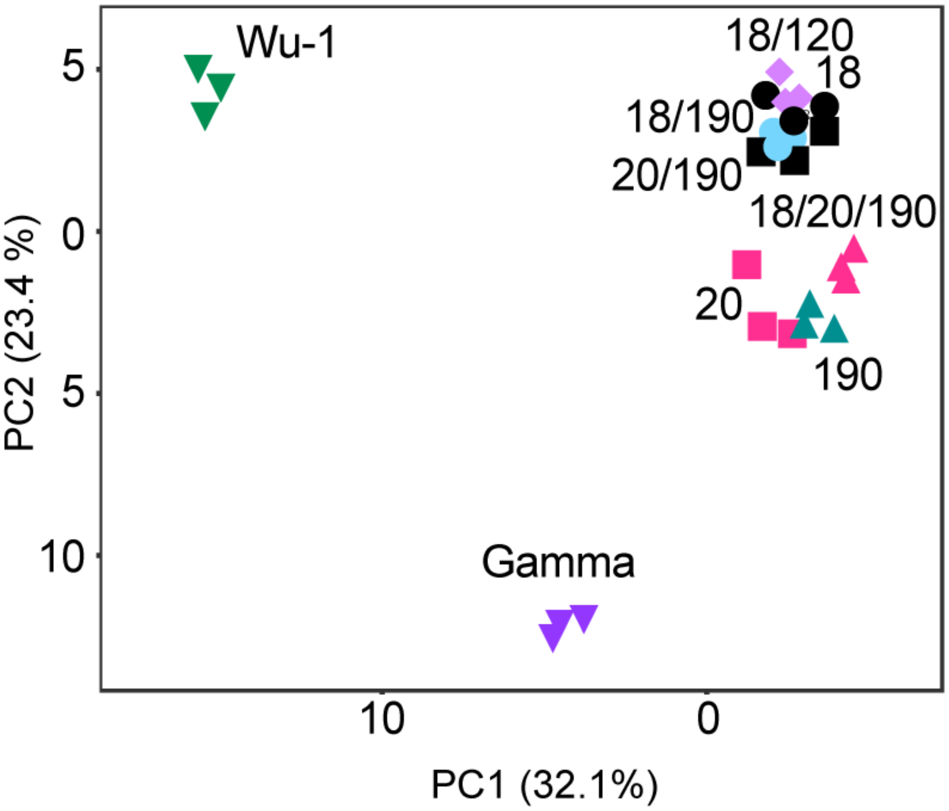


**B. Batch 2**

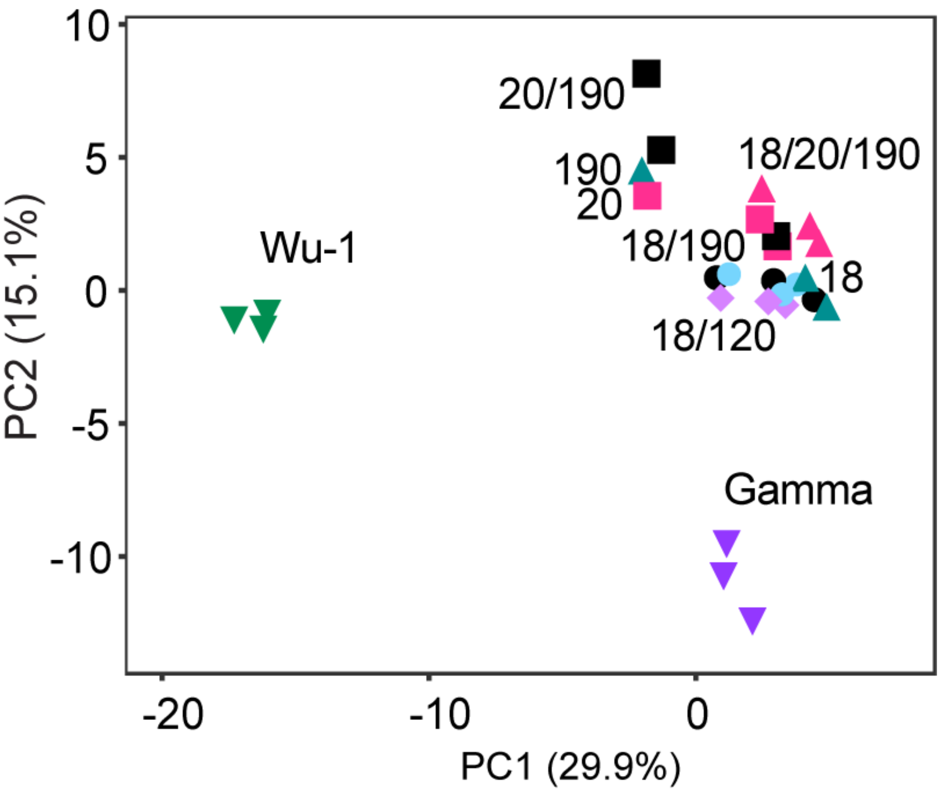

**Supplementary Figure 3.** Clustered heatmap of the relative abundance of all identified glycoforms in all replicates excluding N188. Both rows and columns were clustered using correlation distance and average linkage. The proteins names 18, 20 and 190 represent the proteins with the mutations L18F, T20N and R190S, respectively.

**A. Batch 1**

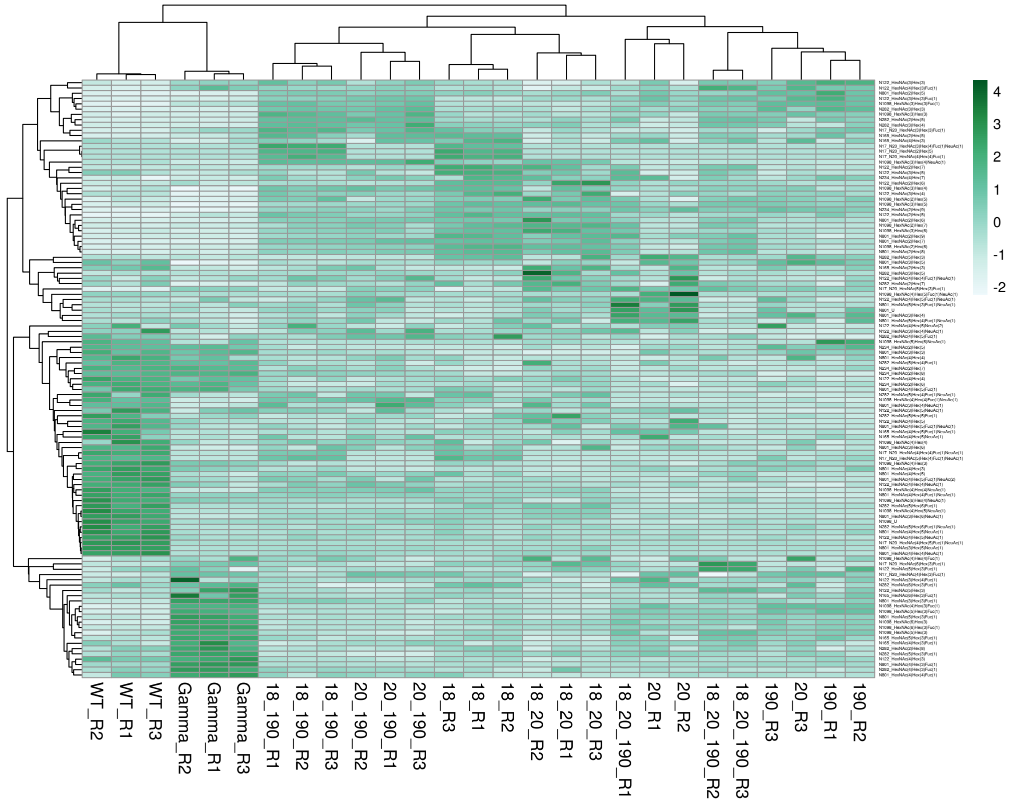


**B. Batch 2**

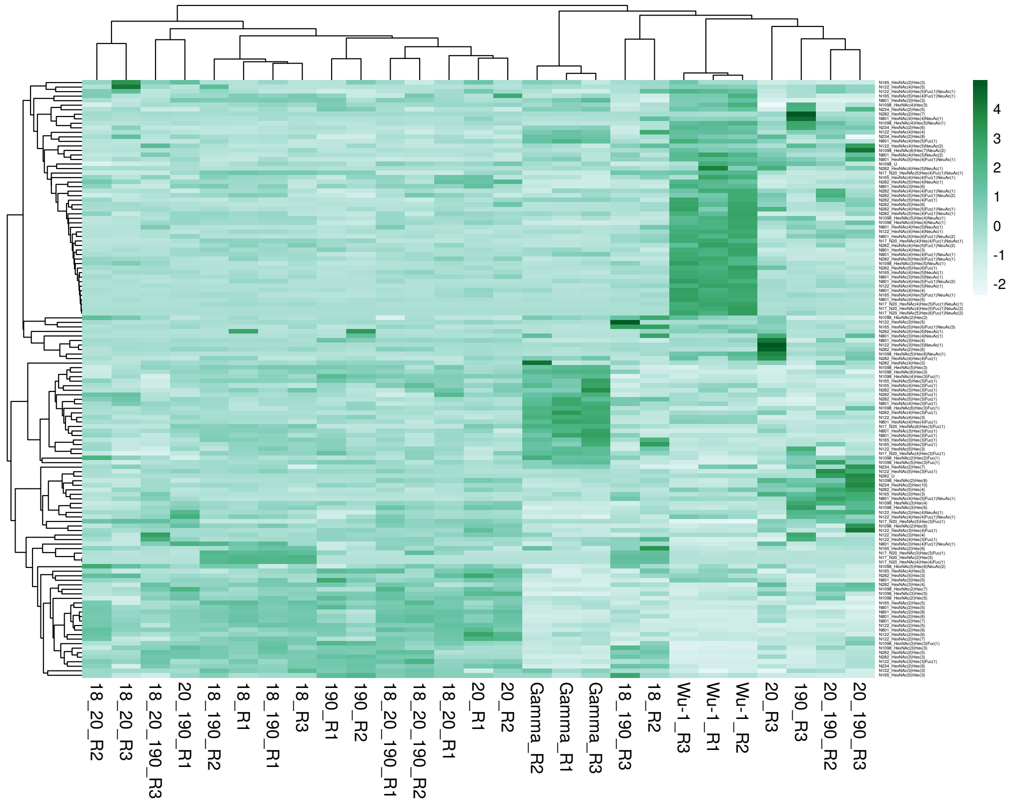
